# Inducible transcriptional condensates drive 3D genome reorganization in the heat shock response

**DOI:** 10.1101/2021.10.27.466170

**Authors:** Surabhi Chowdhary, Amoldeep S. Kainth, Sarah Paracha, David S. Gross, David Pincus

## Abstract

Mammalian developmental and disease-associated genes concentrate large quantities of the transcriptional machinery by forming membrane-less compartments known as transcriptional condensates. However, it is unknown whether these structures are evolutionarily conserved, capable of stress-inducible gene activation or involved in 3D genome reorganization. Here, we identify inducible transcriptional condensates in the yeast heat shock response (HSR). HSR condensates are biophysically dynamic spatiotemporal clusters of the sequence-specific transcription factor Heat shock factor 1 (Hsf1) with Mediator and RNA Pol II. Uniquely, HSR condensates drive the coalescence of multiple Hsf1 target genes, even those located on different chromosomes. Binding of the chaperone Hsp70 to a site on Hsf1 represses clustering, while an intrinsically disordered region on Hsf1 promotes condensate formation and intergenic interactions. Mutation of both Hsf1 determinants reprograms HSR condensates to become mammalian-like: constitutively active without intergenic coalescence. These results suggest that transcriptional condensates are ancient and flexible compartments of eukaryotic gene control.

## INTRODUCTION

Eukaryotes have evolved specialized mechanisms to ensure robust expression of critically important genes such as those activated to specify cell lineage and in response to stress. Mammalian cells utilize clusters of transcription factor binding sites, termed “super-enhancers” (SEs), to drive high-level gene expression by concentrating a large fraction of transcriptional activators and coactivators such as the Mediator complex (Hnisz et al., 2013; Whyte et al., 2013). SEs are interwoven by an extensive DNA looping network constituted by enhancer elements and target gene promoters (Beagrie et al., 2017; Dowen et al., 2014; Huang et al., 2018). Recent studies suggest that SEs function through cooperative assembly of the transcriptional apparatus into “biomolecular condensates” (Boija et al., 2018; Guo et al., 2019; Sabari et al., 2018), defined as self-organized membrane-free compartments that concentrate specific biomolecules (Banani et al., 2017). Transcriptional condensates at SEs are hypothesized to drive transcriptional bursting and stabilize expression of associated genes (Henninger et al., 2021; Hnisz et al., 2017; Sabari et al., 2018).

Biomolecular condensates in general – and transcriptional condensates in particular – are commonly established and maintained by multiple low affinity interactions among their components, often involving intrinsically disordered regions (IDRs) in proteins (Banani et al., 2017; Shin and Brangwynne, 2017). IDRs engage in a network of weak and transient “fuzzy” interactions with other proteins and nucleic acids and are often subject to reversible post-translational modifications that can regulate these interactions (Boehning et al., 2018; Boija et al., 2018; Chong et al., 2018; Guo et al., 2019; Hnisz et al., 2017; Sabari et al., 2018; Sanborn et al., 2021; Shin and Brangwynne, 2017; Tuttle et al., 2021; Wei et al., 2020). Above a critical concentration threshold, the interacting proteins and/or nucleic acids acquire low free energy states by spontaneously separating into two distinct phases: a dense, macromolecule-enriched phase and a dilute, macromolecule-depleted phase (Banani et al., 2017; Shin and Brangwynne, 2017). The nature and strength of intermolecular interactions and the number of participant molecules are likely to influence the dynamics inside the condensates (Brangwynne et al., 2009; Hnisz et al., 2017). In the case of transcriptional condensates, intermediate concentrations of transcriptional activators can lead to formation of active hubs that are not necessarily phase separated (Chong et al., 2018; Baek et al., 2021), suggesting biophysical diversity among different transcriptional condensates.

To date, transcriptional condensates have only been described at stably expressed loci in mammalian cells (Sabari et al., 2020). Whether they regulate activation of highly-expressed inducible genes, such as those activated by environmental stress, and whether they are conserved features of eukaryotic transcription remains unknown. In fact, in the single known example of biomolecular condensation of a stress responsive transcription factor – phase separation of Heat Shock Factor 1 (Hsf1) in human cells – the condensates have been shown to be anti-correlated with transcriptional activation of the heat shock response (HSR) and localized away from canonical HSR genes (Gaglia et al., 2020; Jolly et al., 1997). Moreover, it remains unclear how transcriptional condensates conduct gene control in a three-dimensional (3D) context and whether they influence 3D genomic organization.

Recent studies in budding yeast have revealed profound remodeling of the 3D chromatin architecture following activation of the HSR by Hsf1 (Chowdhary et al., 2017; Chowdhary et al., 2019), and we have hypothesized that transcriptional condensates drive this 3D genomic reorganization (Kainth et al., 2021). Indeed, the HSR shares several key features with mammalian SEs, such as recruitment of high levels of the transcriptional machinery at the associated genes and extensive DNA looping interactions. A key distinction is that, as opposed to the stable gene expression enforced by SEs, HSR activation in yeast is induced in a remarkably rapid yet transient manner: large increases in occupancy of Hsf1 and the transcriptional machinery at HSR genes accompany transcriptional activation within 60 seconds of temperature upshift and return to pre-heat shock levels within 30 to 60 minutes (Chowdhary et al., 2017; Chowdhary et al., 2019; Kim and Gross, 2013; Kremer and Gross, 2009; Pincus et al., 2018; Vinayachandran et al., 2018). Beyond the enhancer-promoter loops found in SEs, HSR genes engage in physical interactions with other HSR genes separated by large distances and even located on different chromosomes (Chowdhary et al., 2017; Chowdhary et al., 2019). Both intragenic and intergenic rearrangements of HSR genes are dynamic and concur with transcriptional output (Chowdhary et al., 2017). Thus, as opposed to the situation described for the HSR in human cells, induction of the yeast HSR may involve transient condensates that form intergenic hubs of transcriptional activity.

Here we provide experimental evidence implicating biomolecular condensation as the underlying mechanism of 3D gene control during the yeast HSR. We show that Hsf1, together with Mediator and RNA Pol II, forms transcriptional condensates at HSR genes in response to heat shock. These condensates are inducible, highly dynamic and promote intergenic chromatin interactions among genes in the Hsf1 regulon. We identify the N-terminal IDR of Hsf1 and a C-terminal binding site for the chaperone Hsp70 as molecular determinants that promote condensation with intergenic interactions and inhibit cluster formation, respectively. Finally, we generate a separation-of-function mutant of Hsf1 by mutating both of these determinants that forms constitutive condensates but fails to drive intergenic interactions. This double mutant displays a strong growth defect at elevated temperature, suggesting that formation of constitutive condensates that lack intergenic interactions comes at a fitness cost in this system. Together, our work reveals that transcriptional condensates are a conserved feature of eukaryotic gene control that can be induced and dissolved as a means to reversibly reconfigure the 3D genome.

## RESULTS

### Inducible high-level occupancy of the transcriptional machinery at HSR genes

To determine whether yeast HSR genes – defined as genes under the control of Hsf1 – share key features with genes that drive lineage specification and disease progression in mammalian cells, we assessed the extent to which HSR genes concentrate the transcriptional machinery. Using a consistent bioinformatic pipeline, we reanalyzed chromatin immunoprecipitation sequencing (ChIP-seq) data of Hsf1, Med15 (subunit of the Mediator complex known to interact with Hsf1) and Rpb1 (subunit of RNA Pol II) across the genome under basal and acute heat shock conditions (Albert et al., 2019; Anandhakumar et al., 2016; Kim and Gross, 2013; Pincus et al., 2018; Sarkar et al., 2020). We found strong enrichment of all three factors at HSR genes upon heat shock and observed that the Hsf1 and Med15 peaks colocalize at heat shock elements (HSEs), the known binding sites for Hsf1 (Figure 1A, B, S1A-C). Rank order analysis of protein-coding genes (n=5983) by change in Med15 occupancy during acute heat shock revealed that all HSR genes are in the top quintile, with the majority residing above inflection point of the curve (Figure 1C). To determine if Hsf1 binding is sufficient to recruit high levels of Mediator upon heat shock, we performed ChIP of Hsf1 and Med15 at the non-HSR gene *BUD3* in a wild type strain and a strain harboring a chromosomal integration of a high-affinity HSE at the *BUD3* locus (Chowdhary et al., 2019). While neither Hsf1 nor Med15 were detectable at the wild type *BUD3* locus, a brief heat shock induced massive recruitment of Hsf1 and Med15 to *BUD3* with the engineered HSE (Figure S1D), suggesting that Mediator is robustly and selectively recruited by Hsf1 at its gene targets. Together, these data demonstrate that HSR genes disproportionately concentrate the transcriptional machinery upon activation.

**Figure 1.**
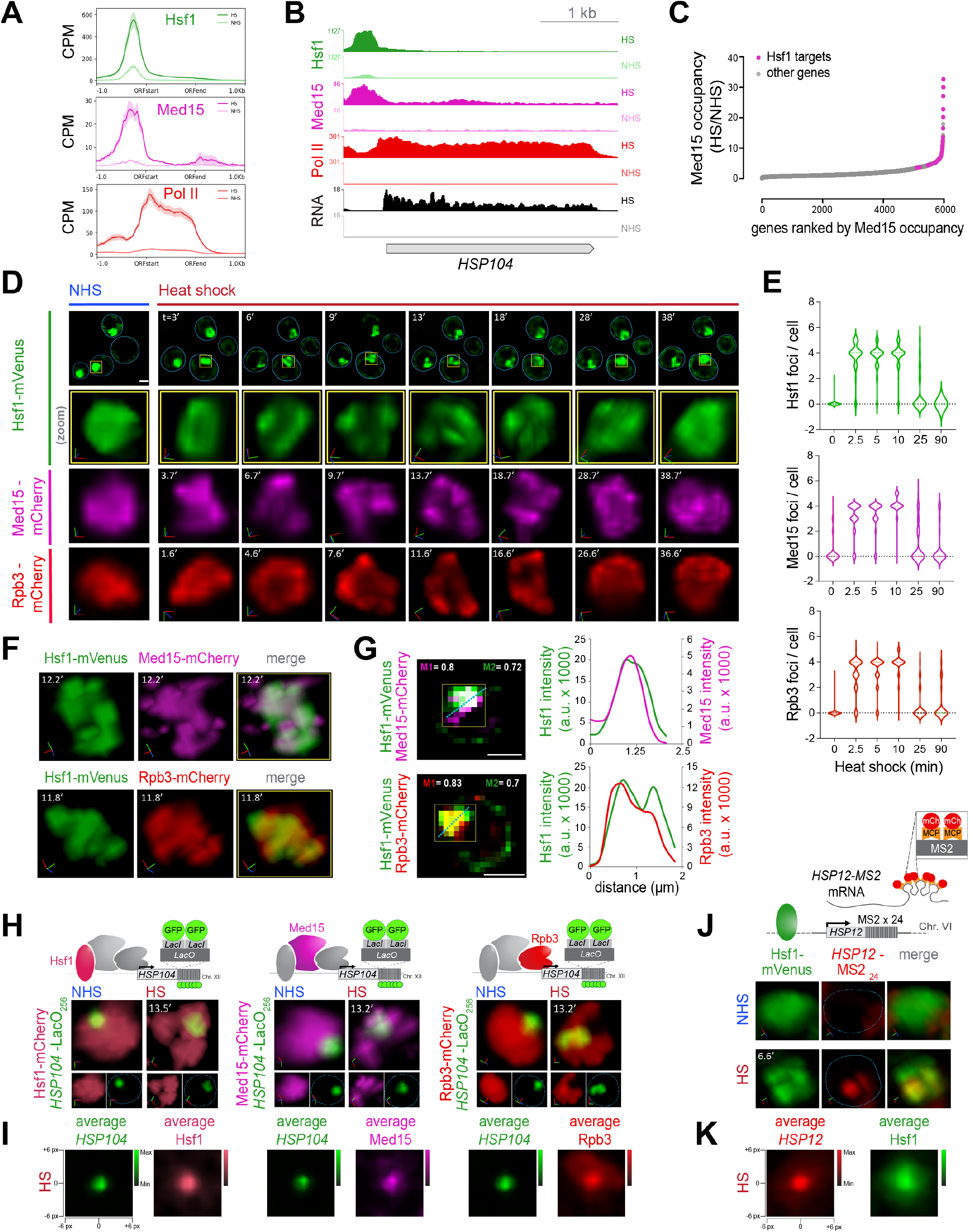
Hsf1, Mediator and RNA Pol II form transcriptionally active clusters at HSR genes upon heat shock. **A)** Metagene plots of Hsf1 (top), Med15 (middle) and Pol II (Rpb1; bottom) normalized ChIP-seq reads of Hsf1-dependent genes in *S. cerevisiae* (n=42) centered over ORFs with 1 kb surrounding region. Shown are means +/- SE (shaded region). NHS, non-heat shock; HS, heat shock; CPM, counts per million reads. **B)** Hsf1, Med15 and Pol II ChIP-seq densities as well as nascent mRNA counts (NAC-seq) at the *HSP104* locus. **C)** Fold change in Med15 occupancy across Pol II-transcribed genes (0 to −1 kb from +1 ATG) upon heat shock, arranged according to their rank order. **D)** Fluorescence imaging analysis of cells expressing Hsf1-mVenus, Med15-mCherry or Rpb3-mCherry before (NHS, 24-26°C) and following HS (39°C) for the times (t) indicated. A single cell is followed throughout the heat shock time course. Row 1: images of cells expressing Hsf1-mVenus. Representative planes are shown. Blue line highlights cell boundary. Scale bar is 2 μm. Row 2: enlarged view of the nucleus (see yellow box in Row 1). Shown is the 3D volumetric rendering of each nucleus; x (red), y (green) and z (blue) axes are indicated. Rows 3 and 4: 3D volumetric rendering of nuclei of cells expressing Med15-mCherry and Rpb3-mCherry, respectively. **E)** Number of Hsf1 (top), Med15 (middle) or Rpb3 (bottom) foci per cell quantified from fixed cell imaging. 50-90 cells were evaluated per time point. **F)** Live imaging of cells co-expressing Hsf1-mVenus and Med15-mCherry (top), or Hsf1-mVenus and Rpb3-mCherry (bottom). Cells were heat-shocked for the indicated time. **G)** Left: Quantification of colocalization using Mander’s Overlay Coefficient of Hsf1-mVenus and Med15-mCherry (top), or Hsf1-mVenus and Rpb3-mCherry (bottom). M1 indicates the fraction of mCherry that overlaps mVenus; M2, fraction overlap of mVenus with mCherry. Scale bar is 2 μm. Right: average intensity profiles along blue dotted lines indicated in the images. **H)** 3D rendered micrographs of Hsf1-mCherry (left), Med15-mCherry (middle) or Rpb3-mCherry (right) and the *HSP104-LacO_256_* gene locus as depicted in the schematics under NHS and following heat shock for the indicated times. Blue dotted line highlights nuclear boundary. **I)** Contour plots of Hsf1-mCherry (left), Med15-mCherry (middle) or Rpb3-mCherry (right) showing averaged signal intensities centered at the *HSP104-LacO_256_* gene locus 13 minutes following heat shock. **J)** Top: schematic of Hsf1 (green) and the MCP-mCherry labeled *HSP12-MS2_24_* mRNA (red). Bottom: 3D rendered micrographs of representative cells heat-shocked for ~7 min or not (NHS). Blue dotted line highlights nuclear boundary. **K)** Contour plot of Hsf1-mVenus (right) showing averaged intensity signal centered at *HSP12-MS2_24_* mRNA (left). Cells were imaged at 5 minutes of heat shock.

### Hsf1, Mediator and RNA Pol II colocalize in subnuclear clusters upon heat shock

We hypothesized that HSR genes engage such a large fraction of the transcriptional machinery by spatially concentrating the requisite factors within the nucleus. Therefore, we investigated the spatiotemporal distributions of Hsf1, Mediator and RNA Pol II in live single cells upon heat shock (Figure 1D). In unstressed cells, endogenously tagged Hsf1-mVenus was predominantly nuclear and diffusely localized. However, Hsf1 formed semi-discrete subnuclear clusters within 6 min of heat shock that persisted for 20-25 min, after which Hsf1 returned to its diffuse localization pattern. To rule out photobleaching artifacts and to quantify how frequently Hsf1 clusters appeared within a population of cells, we performed microscopy of cells that were subjected to instantaneous heat shocks for 0, 2.5, 5, 10, 25 and 90 min and then fixed (Figure S2A). This analysis revealed the presence of Hsf1 clusters in >80% cells subjected to either 2.5, 5, or 10-min heat shock. Consistent with the live imaging, few cells evinced clusters when induced for 25 min or longer (Figure S2B). We typically observed 4 clusters of Hsf1 per individual nucleus, although the number varied between 2 and 5 over the course of heat shock (Figure 1D, E). Analysis of diploid cells expressing Hsf1-GFP gave virtually identical results; neither kinetics nor number of clusters changed significantly, indicating that the number of Hsf1 clusters does not correlate with ploidy (Figure S2C). To distinguish whether redistribution of Hsf1 into clusters is a physical consequence of elevated temperature or a regulatory consequence of Hsf1 activation, we imaged Hsf1-mVenus expressing cells following treatment with azetidine-2-carboxylic acid (AZC), a proline analog that incorporates into nascent proteins, impairs protein folding and activates Hsf1 (Trotter et al., 2001; Zheng et al., 2018). Following AZC treatment, Hsf1 formed intranuclear clusters in a majority of cells (Figure S2D). Therefore, Hsf1 clustering appears to be driven by activation rather than heat per se.

We next tested whether Mediator and RNA Pol II also formed intranuclear clusters during heat shock by imaging haploid cells expressing endogenously-tagged Med15-mCherry or Rpb3-mCherry. In the absence of heat shock, Med15 and Rpb3 were diffusely localized within the nucleus. However, like Hsf1, both formed an average of 4 subnuclear clusters in >80% of cells upon heat shock that emerged and dissolved with similar kinetics to Hsf1 (Figures 1D, E and S2A, B). Moreover, live cell 3D imaging of cells co-expressing Med15-mCherry and Rpb3-mCherry with Hsf1-mVenus revealed that the clusters formed by both factors co-localized with Hsf1 during heat shock with a fractional overlap of at least 70% (Figures 1F, G and S2E). These data provide evidence of spatiotemporal interactions between Hsf1, Mediator and RNA Pol II inside living cells following heat shock.

### Hsf1 clusters activate transcription of HSR genes

In human cells, Hsf1 also forms subnuclear clusters upon activation, but these clusters do not reside at protein coding genes and are therefore not sites of HSR activation (Gaglia et al., 2020; Jolly et al., 1997). To determine whether HSR genes associate with the Hsf1/Med15/Rpb3 clusters in yeast, we created three individual strains bearing 3′-flanking arrays of 256 *lacO*-repeats at the *HSP104* locus – a canonical HSR gene bound by these factors (Figure 1B) – and co-expressing LacI-GFP with either Hsf1-, Med15- or Rpb3-mCherry (Figure 1H). Live-cell imaging revealed that under basal conditions, *HSP104-lacO_256_* minimally overlapped with Hsf1-, Med15- and Rpb3-mCherry but consistently co-localized with these factors during acute heat shock (Figure 1H and S2F). Across a population of cells, the average fluorescence intensities of Hsf1, Med15 and Rpb3 were enriched at the center of the *HSP104-lacO_256_* focus (Figure 1I). These results indicate that the Hsf1, Mediator and Pol II clusters associate with a canonical HSR gene.

To directly test whether Hsf1 clusters are sites of active transcription, we employed the MS2 RNA imaging system to visualize nascent transcript production (Hocine et al., 2013; Lenstra and Larson, 2016). We inserted 24 repeats of the MS2 stem loop into the genome in the 3′ untranslated region of the *HSP12* gene, another HSR target (Figure S1C), in a strain expressing the MS2 coat protein fused to mCherry (MCP-mCherry) and co-expressing Hsf1-mVenus. We observed no HSP12-MS_24_ fluorescence under non-heat shock conditions, indicating a lack of transcription of *HSP12* mRNA, while heat shock induced the transient appearance of a concentrated focus of nascent *HSP12-MS_24_* mRNA that colocalized with Hsf1-mVenus (Figures 1J and S2G). The average fluorescence intensity of Hsf1 was enriched at the center of the *HSP12-MS2_24_* mRNA focus across a population of acutely heat-shocked cells (Figure 1K). These data demonstrate that, upon heat shock, Hsf1 forms subnuclear clusters with the transcriptional machinery that mark sites of nascent mRNA production.

### Hsf1 clusters exhibit properties of biomolecular condensates

As shown above, HSR genes concentrate high levels of the transcriptional machinery and are activated in spatial clusters, reminiscent of SE-associated transcriptional condensates that drive expression of mammalian cell identity genes. Transcriptional condensates display the hallmark property of rapid exchange and reorganization of constituent biomolecules (Boija et al., 2018; Cho et al., 2018; Sabari et al., 2018). To investigate the dynamics of Hsf1 clusters in living cells, we employed super-resolution imaging. We took time-series images of cells expressing Hsf1-mVenus every minute and found that the Hsf1 clusters exhibited a variety of temporal signatures along the course of heat shock: (1) an early demixing phase; (2) an intermediate intermixing phase; and (3) a late remixing phase (Figure 2A). The demixing phase is marked by formation of Hsf1 clusters during the first 6-7 min of heat shock induction. In the intermixing phase (8-27 min), Hsf1 clusters rearrange continuously within the nucleus, demonstrating the exchange of contents among the clusters. In the remixing phase (28-32 min), the contents of the clusters dispersed to the pre-heat shocked diffuse form.

**Figure 2.**
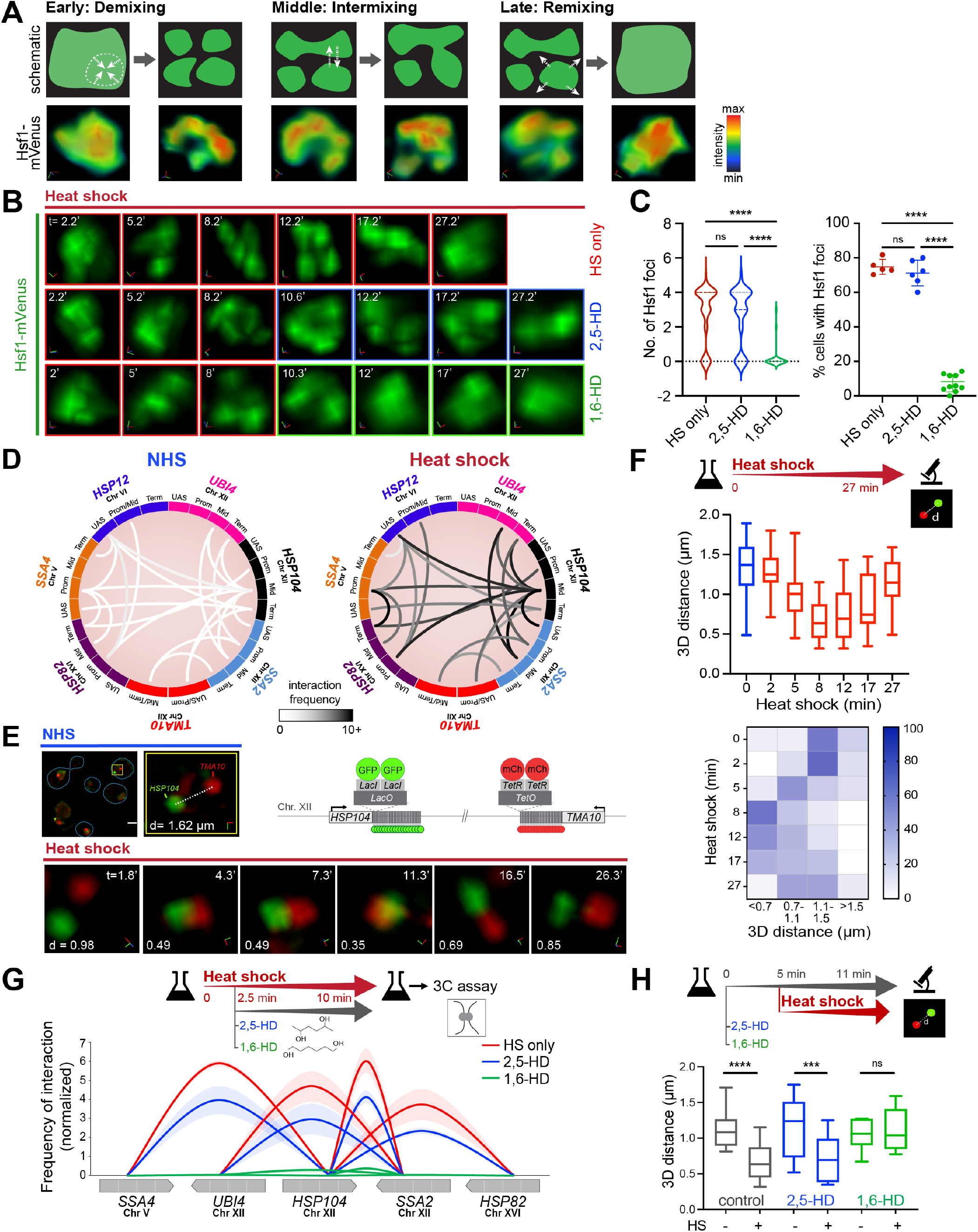
Hsf1 clusters are dynamic and associated with HSR gene coalescence. **A)** Schematic and super-resolution (STED) imaging of Hsf1-mVenus dynamics showing the different stages at heat shock times: 2 and 7 min (early), 8 and 11 min (mid), 28 and 32 min (late). 3D rendering axes: x (red), y (green) and z (blue) are shown. **B)** 3D rendered micrographs of a representative cell subjected to heat shock for times (t) indicated in the absence of alcohol (top) or treated with either 2,5-HD (middle) or 1,6-HD (bottom) after 9 min of HS. Note that a different cell was imaged after alcohol treatment. **C)** Number of Hsf1 foci per cell and the percentage of cells showing >2 Hsf1 foci (n=140-190 cells/condition) following heat shock for 9 minutes +/- the indicated HD. Cells were imaged approximately 3 minutes after adding HD. ****P <0.0001; ns (not significant), P >0.05. P values were calculated by ANOVA followed by Tukey’s post hoc analysis. **D)** Circos plots depicting intergenic interactions between indicated gene pairs as determined by Taq I-3C derived from (Chowdhary et al., 2017; Chowdhary et al., 2019). Arc shades depict the frequencies of chromatin interactions between indicated regions. For each gene pair, top two interactions are shown. **E)** Live-cell imaging of a diploid strain expressing *HSP104-lacO_256’_, TMA70-tetO_200’_, LacI-GFP* and *TetR-mCherry* (schematic, top right). Top left: micrographs of representative cells under NHS conditions (blue line highlights cell boundary) and an enlarged 3D view of the nuclear region indicated within the yellow box. Bottom: enlarged 3D micrographs of a representative cell subjected to heat shock for the times (t) indicated (axes: x (red), y (green) and z (blue)). d, 3D distances measured between signal centroids. **F)** Distribution of 3D distances between *lacO*-tagged *HSP104* and *tetO*-tagged *TMA10* gene loci in cells subjected to heat shock for times indicated (n=13 for NHS; n=20-30 for HS time points). Bottom: Heat map depicting percentage of cells with tagged gene loci within indicated 3D distances binned at intervals of 0.4 μm. **G)** Intergenic contacts (solid arcs) between indicated Hsf1 target gene pairs as determined by TaqI-3C. Values indicate normalized interaction frequencies. Genes are segmented as UAS, promoter, mid-ORF and terminator (from blunt to arrowhead direction). Data are derived from two independent biological replicates; qPCR =4. Depicted are means +/- SD (shaded region around solid arcs). **H)** Distribution of 3D distances between tagged gene loci under conditions as indicated. Cells were left untreated or pre-treated with either 2,5- or 1,6-hexanediol followed by no heat shock or heat shock for 6 min and imaged at times (t) indicated. n =13-22. ****P <0.0001; ***P <0.001; ns (not significant), P >0.05 (calculated using two-tail t test).

To further probe the material properties of Hsf1 clusters, we tested their sensitivity to an aliphatic alcohol, 1,6- hexanediol (1,6-HD). 1,6-HD is widely used to dissolve biomolecular condensates of liquid-like character – including transcriptional condensates in mammalian cells – because it can disrupt weak interactions, particularly those that are hydrophobic in nature (Cho et al., 2018; Kroschwald et al., 2015; Patel et al., 2015; Sabari et al., 2018). Extended treatment of cells with 1,6-HD can impact membrane integrity and cause blebbing – which could be an early sign of autophagy or cell death – as well as disrupt chromatin architecture and inhibit kinase and phosphatase activity (Duster et al., 2021; Itoh et al., 2021; Kroschwald et al., 2017). To minimize the pleiotropic and aberrant effects of 1,6-HD, we optimized the concentration and duration of hexanediol treatment for the two most widely studied examples of nuclear biomolecular condensates: the nucleolus and the nuclear pore complex (NPC) (Banani et al., 2017; Kato and McKnight, 2018). To this end, we applied 1,6-HD to living cells expressing endogenous Nop56-mRFP (nucleolar component) and Pom34-GFP (nuclear pore component) at three different concentrations (3, 5 and 10%) for 20-30 minutes (Figure S3A, B). While it took more than 15 minutes to disintegrate the nucleolus and NPC with 3% 1,6-HD, similar effects were evident in less than 4 minutes when 10% 1,6-HD was applied. Moreover, membrane blebbing was prominent following addition of 10% 1,6-HD. Treatment with 5% 1,6-HD resulted in dissolution of the nucleolus and NPC after 7 minutes with no signs of membrane blebbing until after 20 minutes (Figure S3B). We also scored the effects of 2,5-hexanediol (2,5-HD), a less hydrophobic alcohol with the same atomic composition as 1,6-HD. 2,5-HD had minimal effect on nucleolar and NPC integrity when used at lower concentrations (3 or 5%). However, at 10%, the effects of 2,5-HD were the same as observed with 1,6-HD (Figure S3A, B). Based on these observations, we identified 5% for less than 20 minutes as the optimum concentration and treatment duration for both hexanediols.

To determine the effect of hexanediols on Hsf1 clusters, we first heat shocked cells for 8 minutes to allow cluster formation and then treated cells expressing Hsf1-mVenus with either 5% 1,6-HD or 2,5-HD. Hsf1 clusters were dissolved within 4 minutes upon addition of 1,6-HD but were unaffected by 2,5-HD treatment and mimicked cells that were only heat-shocked (Figure 2B, C). We observed similar effects on Med15 clusters, except they were even more rapidly dissolved upon 1,6-HD treatment (Figure S3C, D). Together, these data indicate that the transcriptionally active Hsf1 clusters display properties characteristic of biomolecular condensates: they continuously intermix and exchange contents, and they are potentially held together by weak, hydrophobic interactions. We will hereon refer to them as HSR condensates.

### HSR condensates drive intergenic interactions among HSR genes

HSR genes engage in selective intra- and inter-chromosomal interactions upon heat shock-induced activation (Chowdhary et al., 2017; Chowdhary et al., 2019), and we hypothesized that these interactions are driven by condensate formation (Kainth et al., 2021). Using a highly sensitive chromosome conformation capture technique (TaqI-3C) (Chowdhary et al., 2020), we have demonstrated interactions among seven Hsf1-dependent genes spanning distinct chromosomes (Chowdhary et al., 2019) (Figure 2D). The intergenic interactions between Hsf1-dependent genes occur with low frequency under non-heat shock conditions but increase many fold upon acute heat shock, coincident with their increased transcription. To complement this molecular assay, we directly imaged intergenic coalescence of HSR genes in live cells by utilizing a heterozygous diploid strain bearing operator arrays chromosomally linked to HSR genes, *HSP104-lacO_256_* and *TMA10-tetO_200’_*, that co-expresses LacI-GFP and TetR-mCherry (Chowdhary et al., 2019) (Figure 2E). Under non-heat shock conditions, *HSP104-lacO_256_* and *TMA10-tetO_200_*were spatially separated, as indicated by a large 3D distance between the two loci (d =1.6 μm). This is consistent with the respective locations of *HSP104* and *TMA10* on the far left and right arms of chromosome XII, as well as their physical isolation from each other due to the nucleolar barrier between them. However, upon heat shock, *HSP104-lacO_256_* and *TMA10-tetO_200_*came in close proximity within 5 minutes of thermal exposure, coalescing into a diffraction-limited single focus after 10 minutes that ultimately dissipated after 16 minutes (Figure 2E). Across a population of cells, *HSP104-lacO_256_* and *TMA10-tetO_200_* were separated by an average 3D distance of <0.7 μm in acutely heat-shocked cells, and >1.1 μm under non-inducing and chronic heat shock conditions (17 min and later) (Figure 2F). These data are consistent with our previous observations of chromosomally unlinked HSR genes (Chowdhary et al., 2017) and indicate that the coalescence of Hsf1-dependent genes is highly dynamic with kinetics closely paralleling that of Hsf1 cluster formation and dissolution.

To determine if HSR condensates are involved in driving intergenic coalescence, we perturbed Hsf1 clusters and assayed for coalescence using the TaqI-3C and live single cell imaging analyses. First, we induced Hsf1 clusters by subjecting cells to brief heat shock treatment for 2.5 minutes before adding 1,6-HD or 2,5-HD and performed TaqI-3C (Figures 2G, S3E). We detected strong chromatin interactions between HSR genes in cells heat-shocked for 2.5 minutes. The 3C interactions were only moderately reduced in the presence of 2,5-HD, including intragenic interactions within the HSR genes. By contrast, addition of 1,6-HD reduced the contact frequencies to background levels. Next, we pre-treated cells with 1,6-HD or 2,5-HD prior to heat shock and performed single cell imaging of the strain bearing fluorescently tagged alleles of *HSP104* and *TMA10*. The coalescence of *HSP104-lacO_256_* and *TMA10-tetO_200_* was abolished in cells that were pretreated for 5 min with 5% 1,6-HD before heat shock but occurred normally in cells treated with 5% 2,5-HD, matching the cells with no alcohol pretreatment (Figure S3F). Quantification revealed that the mean 3D distance decreased from >1.0 μm to <0.7 μm upon heat shock over a population of cells subjected to either 5% 2,5-HD or no pre-treatment, whereas the loci remained separated by >1.0 μm in 1,6-HD-treated cells (Figure 2H). These results indicate that 1,6-HD not only dissolves the clusters of Hsf1, but also disrupts intergenic interactions between HSR genes.

### Hsp70 binding represses Hsf1 cluster formation

Do the molecular mechanisms that control Hsf1 activity also regulate HSR condensate formation? Hsf1 activity is repressed when it is bound by the chaperone Hsp70 (Figure 3A) (Krakowiak et al., 2018; Masser et al., 2019; Peffer et al., 2019; Zheng et al., 2016), so we wondered if Hsp70 binding would also repress HSR condensate formation. To test this, we disrupted the Hsp70 binding site located in the C-terminal segment of Hsf1, known as “conserved element 2” (CE2), by introducing alanine substitutions at residues known to be required for Hsp70 association (Krakowiak et al., 2018; Peffer et al., 2019) (Figure 3A, B). We expressed GFP-tagged Hsf1-ce2AAA as the only copy of Hsf1 and imaged its localization. In contrast to the diffuse nuclear localization of wild type Hsf1, we observed constitutive clustering of Hsf1-ce2AAA in unstressed cells that persisted during heat shock (Figure 3C).

**Figure 3.**
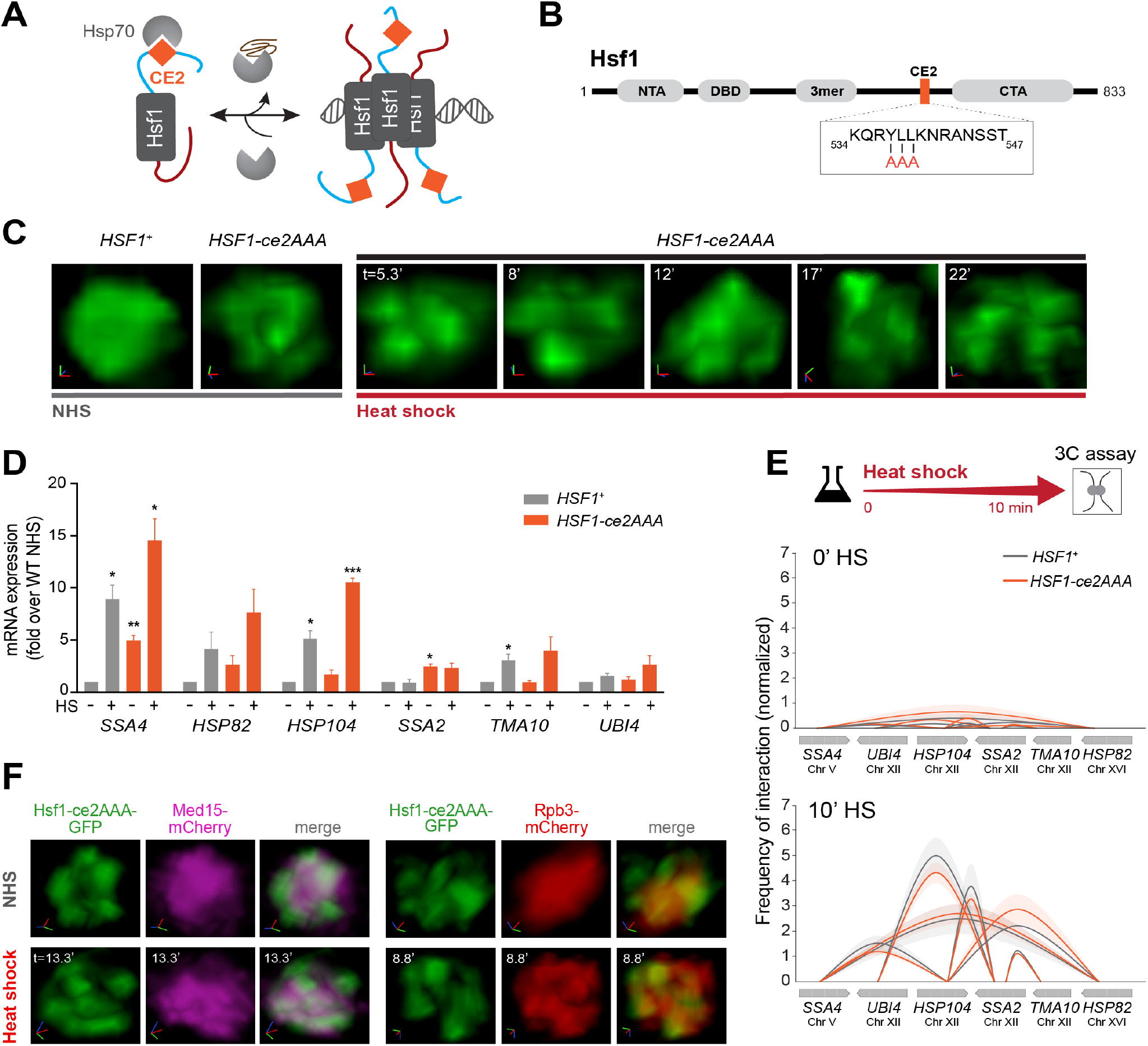
Hsp70 binding to Hsf1 represses cluster formation. **A)** Schematic of the Hsp70 titration model of Hsf1 regulation. Hsp70 binds to Hsf1 via the CE2 domain and inactivates Hsf1. Accumulation of chaperone clients sequesters Hsp70, leaving Hsf1 free to activate transcription. **B)** Domain map of Hsf1. NTA, N-terminal activation domain; DBD, DNA binding domain; 3mer, trimerization domain; CE2, conserved element 2 domain (orange); CTA, C-terminal activation domain. Box: CE2 sequence and alanine substitutions in the Hsf1-ce2AAA mutant. **C)** 3D rendered images of wild type and Hsf1-ce2AAA mutant cells in NHS and HS conditions. D) mRNA expression measured by RT-qPCR of representative Hsf1-regulated genes in *HSF1^+^* and *HSF1-ce2AAA* strains under NHS and 10 min-HS conditions. ***P <0.001; **P <0.01; *P <0.05 (calculated using multiple unpaired t tests) **E)** Intergenic contacts (solid arcs) between indicated Hsf1 target gene pairs under NHS and 10 min-HS conditions as determined by TaqI-3C. **F)** Live imaging of cells expressing Hsf1 ce2AAA-GFP and Med15-mCherry (left), or Hsf1 ce2AAA-GFP and Rpb3-mCherry (right) under NHS or HS conditions.

To determine whether the constitutive clusters formed by Hsf1-ce2AAA are transcriptionally active, we measured mRNA levels of representative HSR genes in cells expressing wild type Hsf1 and Hsf1-ce2AAA (Figure 3D). In agreement with the known repressive role for Hsp70 binding, we detected a significant increase in the basal mRNA levels of some targets in Hsf1-ce2AAA cells compared to cells expressing wild type Hsf1. However, the basal levels of most target transcripts in the mutant were lower than their respective levels in wild type cells following heat shock, suggesting that the constitutive clusters formed by Hsf1-ce2AAA are less active than those formed by wild type Hsf1 upon activation. Further supporting a lack of full activation in the mutant, both Med15 and Rpb3 were diffusely localized under basal conditions in Hsf1-ce2AAA cells and only formed clusters with Hsf1-ce2AAA upon heat shock (Figure 3F). Moreover, HSR genes showed no detectable intergenic interactions as measured by TaqI-3C under basal conditions in Hsf1-ce2AAA cells, but both intra- and intergenic interactions were induced upon heat shock to the same degree as in wild type cells (Figures 3E and S4B). Thus, mutation of the CE2 binding site for Hsp70 results in constitutive Hsf1 cluster formation, but this is not sufficient to trigger formation of HSR condensates that drive intergenic interactions.

### The N-terminal domain of Hsf1 promotes formation of HSR condensates

Hsf1 contains large functional N-terminal and C-terminal activation domains referred to as NTA and CTA (Figure 4A). The NTA both represses Hsf1 under basal conditions – likely due to a second Hsp70 binding site – and plays a positive role in transactivation, while the CTA simply constitutes a strong transactivation domain (Kim and Gross, 2013; Krakowiak et al., 2018; Peffer et al., 2019; Sorger, 1990). Both regions are enriched in acidic amino acid residues, are highly disordered and facilitate interactions between Hsf1 and Mediator (Kim and Gross, 2013; Peteranderl et al., 1999) (Figures 4B and S4C). To assess their roles in intergenic interactions and HSR condensate formation, we separately removed both regions.

**Figure 4.**
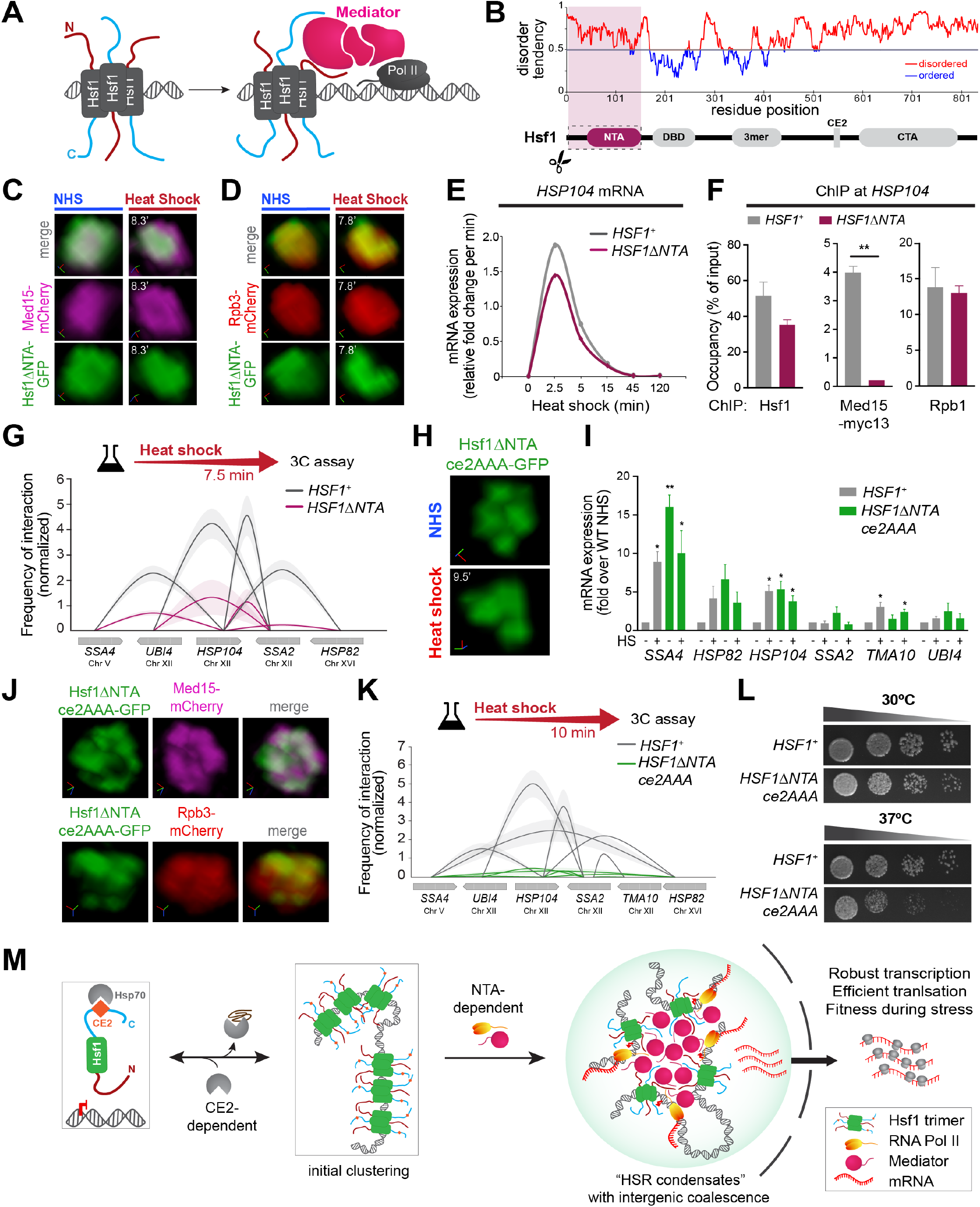
HSR condensates drive inducible intergenic coalescence by the combined actions of the CE2 and NTA regions of Hsf1. **A)** Cartoon depiction of Mediator recruitment at HSR genes via dual Hsf1 activation domains. **B)** Domain map and prediction of disorder in Hsf1 by IUPRED. **C)** Live imaging of cells expressing Hsf1ΔNTA-GFP and Med15-mCherry under NHS or HS conditions. **D)** Live imaging of cells expressing Hsf1ΔNTA-GFP and Rpb3-mCherry under NHS or HS conditions. **E)** Transcription rates of *HSP104* in *HSF1^+^* and *HSF1-ΔNTA* isogenic strains during a heat shock time course, as deduced from RT-qPCR. **F)** Hsf1, Med15-myc13 and Rpb1 ChIP analysis at the *HSP104* locus in wild type and *Hsf1-ΔNTA* strains heat-shocked for 7.5 min. Depicted are means + SD. **, P <0.01 (calculated using two-tail t-test). **G)** Intergenic contacts (solid arcs) between indicated Hsf1 target gene pairs in 7.5 min-HS conditions as determined by TaqI-3C. **H)** Representative 3D rendered micrographs of cells expressing Hsf1-GFP and Hsf1-ΔNTA-ce2AAA-GFP under NHS and HS (axes: x (red), y (green) and z (blue)). **I)** mRNA expression measured by RT-qPCR of representative Hsf1-regulated genes in *HSF1^+^* and *HSF1-ΔNTA-ce2AAA* strains under NHS and 10 min-HS conditions. **P <0.01; *P <0.05 (calculated using multiple unpaired t tests) **J)** Live imaging of cells co-expressing Hsf1ΔNTAce2AAA-GFP with Med15-mCherry (top) or Rpb3-mCherry (bottom) under NHS conditions. **K)** Intergenic contacts (solid arcs) between indicated Hsf1 target gene pairs following 10 min-HS conditions as determined by TaqI-3C. **L)** Spot dilution analysis of *HSF1^+^* and *HSF1-ΔNTAce2AAA* cells. Plates were incubated at 30°C or 37°C for 30 h. **M)** Schematic depiction of HSR condensation. In unstressed cells, Hsp70 binds to Hsf1 via the CE2 domain and inactivates Hsf1. Accumulation of misfolded proteins sequesters Hsp70 and frees Hsf1 which then binds to its gene targets and forms initial clusters. The activated Hsf1 recruits Mediator via the NTA domain. Hsf1, together with Mediator and RNA Pol II, forms dynamic transcriptional condensates (HSR condensates) that localize multiple Hsf1 target genes engaging in specific interactions with each other. The HSR condensates with intergenic coalescence promote high transcriptional activity, maximal gene expression and cellular fitness during stress.

Since the NTA represses Hsf1 basal activity, we expected its removal would mimic mutation of the CE2, resulting in constitutive clustering. However, Hsf1-ΔNTA-GFP failed to form clusters even upon heat shock (Figure 4C, D). Moreover, Rpb3 and Med15 also could not re-distribute into clusters in the Hsf1-ΔNTA mutant, suggesting that interaction of Hsf1 with Mediator and Pol II is required for clustering during heat shock (Figure 4C, D). These data further support the interpretation that clustering of Hsf1, and additionally Med15 and Rpb3, is driven by regulatory activation rather than heat per se. At the molecular level, we found that while Hsf1-ΔNTA was able to robustly bind to the promoter of the target gene *HSP104* and recruit RNA Pol II upon heat shock, it was unable to recruit high levels of Med15 (Figure 4F). Since the rate of inducible transcription of *HSP104*, and consistently the intragenic interactions within HSR genes (Figure S4A), was only modestly reduced in the Hsf1-ΔNTA cells relative to wild type (Figure 4E), and Mediator is indispensable for transcriptional activation (Anandhakumar et al., 2016), the lack of detectable Med15 in the ChIP assay suggests that Hsf1-ΔNTA is only able to transiently recruit Mediator but not form a stable interaction. Along with the lack of Med15 recruitment, we observed a substantial decrease in intergenic interactions between *HSP104* and other HSR genes in Hsf1-ΔNTA cells compared to wild type (Figure 4G). Loss of the CTA likewise decreased Mediator recruitment and ablated heat shock-induced intergenic interactions among HSR genes, but it also impaired fitness under all growth conditions, rendering it too severe a mutation to interpret specifically (Figure S4C-F) (Krakowiak et al., 2018; Sorger, 1990) These data indicate that the NTA promotes the ability of Hsf1 to cluster with Mediator and Pol II upon heat shock and drive intergenic interactions among HSR genes.

### Reprogramming Hsf1 to mimic mammalian transcriptional condensates

HSR condensates share biophysical properties and molecular constituents with mammalian transcriptional condensates that form at SEs, yet HSR condensates are distinct in their inducible and transient kinetics and their ability to drive intergenic interactions. Since loss of CE2 resulted in constitutive Hsf1 clustering and partial de-repression of Hsf1 transcriptional activity, and loss of the NTA reduced intergenic interactions and also partially de-repressed Hsf1, we reasoned that combining these mutations would fully de-repress Hsf1 and reprogram Hsf1 to be mammalian-like: constitutively clustered but lacking intergenic interactions. To test this, we generated a double mutant, Hsf1-ΔNTAce2AAA, GFP-tagged it and expressed it as the only copy of Hsf1 from the endogenous *HSF1* promoter. Indeed, Hsf1-ΔNTAce2AAA-GFP formed constitutive clusters and was fully de-repressed, driving transcription of some target genes under basal conditions to levels exceeding their expression during heat shock in wild type cells (Figure 4H, I). In contrast to the constitutive clusters we observed in the Hsf1-ce2AAA single mutant, Hsf1-ΔNTAce2AAA-GFP colocalized in clusters with Med15- and Rpb3-mCherry under basal conditions (Figure 4J). Despite this high transcriptional activity and clustering, Hsf1-ΔNTAce2AAA was unable to drive intergenic interactions among HSR genes, even during heat shock (Figure 4K). Importantly, Hsf1-ΔNTAce2AAA cells showed impaired growth, especially at elevated temperature, suggesting that the ability to form inducible intergenic transcriptional condensates promotes fitness during stress (Figure 4L). Thus, the CE2 and NTA are determinants that endow HSR condensates with inducibility and the capacity to drive intergenic interactions, features that distinguish them from the transcriptional condensates described at SEs in mammalian cells (Figure 4M).

## DISCUSSION

Here we have shown that activation of the HSR in budding yeast involves the formation of inducible biomolecular condensates that localize multiple target genes into subnuclear compartments with high transcriptional activity. These HSR condensates form and dissolve over the course of heat shock and contain biophysically dynamic clusters of the transcription factor Hsf1 along with Mediator and RNA Pol II (Figure 4M). Our discovery of HSR condensates in yeast demonstrates that transcriptional condensates exist outside of the mammalian lineage and suggests that these structures are likely an ancient and conserved feature of eukaryotic gene control.

As originally described at mammalian SEs, transcriptional condensates are stable structures that activate high-level transcription of individual genes (Boija et al., 2018; Cho et al., 2018; Sabari et al., 2018). The novel features of HSR condensates – their inducible kinetics and intergenic interactions – indicate that transcriptional condensates are capable of rapidly and reversibly remodeling the 3D genome in response to environmental cues. HSR condensates encode these unique properties within the primary sequence of Hsf1 via Hsp70 binding sites and the intrinsically disordered NTA. As such, genetic disruption of the CE2 binding site for Hsp70 combined with ablation of the NTA was sufficient to reprogram HSR condensates to be constitutive structures that lack intergenic coalescence, recapitulating the characteristics of mammalian SE condensates. Thus, relatively simple genetic changes can have dramatic consequences in the regulatory properties of transcriptional condensates, endowing these structures with evolutionary plasticity.

In many biomolecular condensates, especially those forming at gene regulatory elements, the constituent biomolecules contain intrinsically disordered regions (IDRs) that assemble cooperatively (Banani et al., 2017; Boehning et al., 2018; Boija et al., 2018; Chong et al., 2018; Hnisz et al., 2017; Nair et al., 2019; Sabari et al., 2018; Shin and Brangwynne, 2017). Given that deletion of the NTA domain of Hsf1 results in loss of condensates and intergenic coalescence (Figure 4C, D, G), it is likely that cooperative binding among the IDRs of Hsf1, Mediator and RNA Pol II (Figure S4G, H) could serve to nucleate the HSR condensates and bring HSR genes in 3D proximity (Figure 4M). The other possibility is that Hsf1 initiates the condensation. This initiation mechanism could entail cooperative binding of Hsf1 trimers to multiple HSEs of canonical gene targets (Erkine et al., 1999), with each activator molecule then tethering or engaging one or more polymerases and/or Mediator subunits (Baek et al., 2021). These local multivalent interactions between Hsf1 and Mediator at individual HSR genes could then intermix with other such local assemblies leading to intergenic coalescence.

The role of the chaperone Hsp70 in repressing Hsf1 clustering represents a cell biological manifestation of the known role of Hsp70 in repressing Hsf1 transcriptional activity (Kmiecik et al., 2020; Krakowiak et al., 2018; Masser et al., 2019; Zheng et al., 2016). While we do not elucidate the biochemical basis of this repression in the current study, two mechanisms are possible: Hsp70 could passively block clustering by steric occlusion, and/or Hsp70 could mechanically pull Hsf1 out of the clusters. While precedents exist for both of these putative mechanisms in repression of cytosolic condensates (Ruff et al., 2021; Yoo et al., 2021), this role for Hsp70 in regulating Hsf1 clustering is the first time chaperones have been implicated in regulating transcriptional condensates. Regardless of the specific mechanisms, Hsp70 may generally function as a “molecular stir bar” to antagonize condensates both within and outside the nucleus. Expanding on this notion, Hsf1 may have evolved to form condensates upon activation that mimic other condensates recognized by Hsp70 in order to peg its activity – and therefore the expression of the HSR regulon – to the availability of Hsp70. Other chaperones such as Hsp90, which is known to associate with nuclear hormone receptors that also condense with the transcriptional machinery (Frank et al., 2021; Nair et al., 2019; Picard et al., 1990), may also remodel transcriptional condensates.

Are HSR condensates likely to be conserved in mammalian cells? In human cells, HSF1 does indeed form large stress-induced subnuclear foci (Jolly et al., 1997). However, rather than driving transcription of HSR genes, these foci have been shown to localize to noncoding satellite DNA regions and are anti-correlated with HSR gene expression (Gaglia et al., 2020; Jolly et al., 2004). Despite this evidence, it is also clear that mammalian HSR genes are highly induced following heat shock, suggesting that HSF1 is able to recruit large quantities of the transcriptional machinery (Kainth et al., 2021; Mahat et al., 2016). Thus, in addition to the large HSF1 foci localized to noncoding regions, more advanced imaging techniques may reveal smaller HSF1 foci that condense with Mediator at HSR genes to activate transcription. Moreover, although the organization of mammalian genomes into topologically associated domains (TADs) (Dixon et al., 2012) would seem to preclude the ability of HSR genes to coalesce upon activation like they do in yeast, multiple HSR genes are clustered adjacent to each other in the human genome, potentially allowing intergenic interactions among HSR genes to occur (Kainth et al., 2021). Thus, it is possible that inducible intergenic transcriptional condensates are conserved features of the HSR across the eukaryotic lineage.

Finally, HSR condensates may provide several advantages to cells under stress (Figure 4M). First, condensation serves to concentrate large quantities of the transcriptional machinery to enable robust induction of Hsf1-regulated genes critical to counteracting the stress. Second, the coalescence of multiple target genes separated by large distances on and between chromosomes could reduce search time for repeated transcriptional firing events and coordinate transcriptional bursting across the HSR regulon. Third, spatially coordinated transcription of multiple genes could promote mRNA clustering into “transperons” that may bypass normal mRNA quality control processes and increase efficiency of post-transcriptional steps of gene expression such as nuclear export (Nair et al., 2021; Zander et al., 2016). Fourth, HSR condensates may enable target messages to avoid cytosolic stress granules and thereby promote their privileged translation (Zid and O’Shea, 2014), perhaps by recruiting translation factors or mRNA modifying enzymes. HSR condensates represent the founding example of inducible intergenic transcriptional clusters, but given these potential advantages, it may be the case that analogous structures have evolved to execute other stimulus-regulated transcriptional programs within budding yeast and beyond.

## METHOD DETAILS

### Yeast Strains

For tagging Hsf1, Med15 and Rpb3 with mCherry, PCR amplicons with mCherry coding sequence and homology to 3’-ends of either *HSF1, MED15* and *RPB3* were amplified from plasmid pFA6a-mCherry-hphMX6. The amplicons were transformed into the respective strains for in-frame insertion of mCherry. Primers used in strain construction are listed in Table S6.

For Myc x 9 tagging of Med15, genomic DNA of a previously myc9-tagged *MED15* strain (ASK201) (Anandhakumar et al., 2016) was used as template to amplify MYCx 9::TRP1 cassette flanked by DNA homologous to 3’-end of *MED15*. This amplicon was transformed in strains DSG144 and LRY003 to obtain strains ASK213 and ASK214, respectively. LRY003 is a derivative of previously described strain ASK804 (Chowdhary et al., 2019) in which TRP1 was deleted by replacing with KAN-MX.

For Myc x 13 tagging of Med15, plasmid pFA6a-13Myc-His3MX was used as template to obtain MYC13-HIS3 amplicon with homology to 3’-end of *MED15*. This amplicon was transformed in strains DPY144, DPY417 and DPY418 to obtain strains ASK215, ASK216 and ASK217, respectively.

For MS2-MCP labelling of *HSP12* mRNA, MCP-mCherry-URA3 cassette was amplified from plasmid pSH100 with primers Fw MCP-mCherry-Ura3 and Rv MCP-mCherry-Ura3. This cassette was inserted at the endogenous *URA3* locus in strain DPY032 to obtain strain SCY008. Next, 24xMS2-loxP-KANMX6-loxP cassette was amplified from plasmid pDZ415 using primers Fw HSP12-MS2-loxp-KanMX6-loxp and Rv HSP12-MS2-loxp-KanMX6-loxp. This cassette was inserted in the 3’-UTR region of *HSP12* (immediately after stop codon) of strain SCY008 to obtain strain SCY009. Finally, plasmid pSH69 was transformed to express Cre recombinase in strain SCY010 that led to the removal of KANMX in strain SCY011.

The diploid strain ASK741 was created by crossing a MATα derivative of strain DSG200 (Chowdhary et al., 2019) with MATa DSG200.

Plasmids pNH604-HSF1pr-HSF1-GFP and pNH604-HSF1pr-HSF1(147-833)-GFP were used as templates for quick change PCR (Primers, Fw subCE2-AAA and Rv subCE2-AAA) to create CE2->AAA mutation in WT and ΔNTA HSF1, respectively. These plasmids were linearized with Pme I (New England Biolabs) and transformed in strain DPY034 for integration at the *TRP1* locus to obtain strains DPY1805 and DPY1818. Loss of parental HSF1 plasmid was confirmed by growth on 5-FOA media.

A complete list of strains is provided in Table S1. PCR primer sequences are provided in Table S6.

### Culture Conditions

For microscopy, cells were grown at 30°C in SDC (synthetic dextrose complete) media to early-log density (A_600_ = 0.4-0.5).

For 3C, ChIP and RT-qPCR analyses, cells were grown at 30°C in YPD (yeast extract-peptone-dextrose) to a mid-log density (A_600_ = 0.65-0.8). A portion of the culture was maintained at 30°C as non-heat-shocked (NHS) sample while the remainder (heat-shocked (HS) sample) was subjected to an instantaneous 30°C to 39°C thermal upshift for the indicated duration.

For spot dilution analysis, cells were grown to stationary phase in YPD media. Master suspensions were prepared by diluting the saturated cultures to a uniform cell density (*A*_600_=0.3) and were transferred to a 96-well microtiter plate. These were then serially diluted (five-fold). 4 μl of each dilution were transferred onto solid YPDA plates. Cells were grown at either 30° or 37°C for 30 h.

### Chromosome Conformation Capture

TaqI-3C was conducted essentially as previously described (Chowdhary et al., 2017; 2020; Chowdhary et al., 2019). Briefly, cells were cultured to a mid-log density (A_600_ = 0.8) at 30°C. They were either maintained at 30°C or heat-shocked at 39°C for 10 min (or as indicated), and then crosslinked with formaldehyde (1% final concentration). For 3C assay involving hexanediol treatment, cells were heat-shocked at 39°C for 2.5 min followed by treatment with either 2,5- or 1,6-hexanediol (5% final concentration), and then crosslinked with formaldehyde. Crosslinked cells were harvested and lysed in FA lysis buffer (50 mM HEPES pH 7.9, 140 mM NaCl, 1% Triton X-100, 0.1% Sodium deoxycholate, 1 mM EDTA, 1 mM PMSF) for two cycles (20 min each) of vortexing at 4°C. A 10% fraction of the chromatin lysate was digested using 200 U of Taq I (New England Biolabs) at 60°C for 7 h. Taq I was heat-inactivated at 80°C for 20 min. The digested chromatin fragments were centrifuged, and the pellet was resuspended in 100 μl of 10 mM Tris-HCl (pH 7.5). The Taq I-digested chromatin was diluted 7-fold to which 10,000 cohesive end units of Quick T4 DNA ligase (New England Biolabs) were added. Proximity ligation was performed at 25°C for 2h. The ligated sample was digested with RNase at 37°C for 20 min and then Proteinase K at 65°C for 12 h. The 3C DNA template was extracted using phenol-chloroform and then precipitated.

The interaction frequencies were quantified using qPCR. Quantitative PCR was performed on a CFX Real-Time PCR system (Bio-Rad) using Power SYBR Green PCR master mix (Fisher Scientific). Sequences of 3C primers used in this study are provided in Table S2. Normalization controls were used to account for the following: (i) variation in primer pair efficiencies; (ii) primer dimer background; (iii) variation in the recovery of 3C templates; and (iv) to ensure a ligation-dependent 3C signal. For detailed algorithms to calculate normalized 3C interaction frequencies, see below and (Chowdhary et al., 2020).

### Chromatin Immunoprecipitation (ChIP)

ChIP was conducted essentially as previously described (Chowdhary et al., 2019). Briefly, the cells were heat-shocked at 39°C for 7.5 min (or maintained at 30°C) and crosslinked with 1% formaldehyde. Cells were then harvested and subjected to glass bead lysis in lysis buffer (50 mM HEPES pH 7.5, 140 mM NaCl, 1% Triton X-100, 0.1% Sodium deoxycholate, 1 mM EDTA, 2 mM PMSF, and 250 μg/ml cOmplete™, EDTA-free Protease Inhibitor Cocktail) for 30 min at 4°C. The chromatin lysate was sonicated to an average size of ~250 bp using 40 cycles of sonication (30 sec on/off High-Power setting; Diagenode Biorupter Plus). A 20% fraction of the sonicated chromatin was incubated with one of the following antibodies: 1 μl of anti-Rpb1; 1 μl of anti-Hsf1 (Chowdhary et al., 2019) or 2.5 μl of anti-Myc (Santa Cruz Biotechnology) for 16 h at 4°C. Antibody-chromatin complexes were immobilized on Protein A-Sepharose beads (GE Healthcare) for 16 h at 4°C, then washed sequentially with lysis buffer, high salt buffer (50 mM HEPES pH 7.5, 500 mM NaCl, 1% Triton X-100, 0.1% Sodium Deoxycholate, 1 mM EDTA), wash buffer (10 mM Tris pH 8.0, 250 mM LiCl, 0.5% NP-40, 0.5% Sodium Deoxycholate, 1 mM EDTA) and finally 1x TE (10 mM Tris-HCl pH 8.0, 0.5 mM EDTA). Chromatin was eluted by incubating the beads in elution buffer (50 mM Tris pH 8.0, 1% SDS, 10 mM EDTA) at 65°C for 30 min. RNA and proteins were removed by DNase-free RNase (final concentration of 200 μg/ml; incubation at 37°C for 1 h) and Proteinase K (final concentration of 50 μg/ml; incubation at 60°C for 16 h). The ChIP template was extracted using phenol-chloroform and precipitated in presence of ethanol.

ChIP occupancy signals were quantified using qPCR. Sequences of ChIP primers used in this study are provided in Table S4. The ChIP DNA quantities were deduced from interpolation of a standard curve generated using genomic DNA template. The qPCR signal for each primer combination was normalized to the corresponding signal from the input DNA. The input DNA control was incorporated to correct for variation in the recovery of ChIP DNA templates.

### Reverse Transcription-qPCR (RT-qPCR)

Cells were cultured to a mid-log density (A_600_ = 0.8) at 30°C, and were either maintained at 30°C or heat-shocked at 39°C for times indicated. 20 mM sodium azide was added at appropriate times to terminate transcription. Cells were then harvested and subjected to glass bead lysis in presence of TRIzol (Invitrogen) and chloroform for 10 min at 4°C. Total RNA was precipitated in ethanol. A fraction of total RNA (~20 μg) was treated with DNase I (RNase-free; New England Biolabs) at 37°C for 15 min. DNase I was heat-inactivated at 75°C for 10 min. RNA was purified using the RNA clean and concentrator kit (Zymo Research). 2-3 μg of the purified RNA template and random hexamers were used for preparing cDNA (Superscript IV first-strand synthesis system; Invitrogen).

The cDNA reaction mix was diluted 2-fold, and 2 μl of the diluted cDNA template was used for qPCR reaction. Sequences of primers used for RT-qPCR analysis in this study are provided in Table S5. Relative cDNA levels were quantified using the ΔΔCt method (see Chowdhary et al., 2017). qPCR signal from *SCR1* Pol III transcript was used as a normalization control. This accounted for variation in the recovery of cDNA templates. Relative fold change per minute in mRNA expression was calculated by dividing mean mRNA levels (derived from two independent biological samples) for a given time point by previous time point, and then by the time elapsed in minutes.

### Fluorescence microscopy

For live-cell imaging, cells were grown at 30°C to early log phase (A_600_ = 0.5) in synthetic dextrose complete (SDC) medium. An aliquot of living cells was immobilized onto concanavalin A-coated glass bottom dish. Fresh SDC medium was added in the dish before imaging. Images were taken on Leica TCS SP5 II STED-CW laser scanning confocal microscope (Leica Microsystems, Inc., Buffalo Grove, IL) equipped with GaAsP hybrid detector. Samples were subjected to heat shock (39**°**C) by heating the objective using Bioptechs objective heater system and by controlling the temperature of incubator chamber enclosing the microscope. For experiments involving hexanediol treatment, cells were heat-shocked and imaged for ~8 min. Immediately after this, the SDC media in the dish was replaced with pre-warmed (39**°**C) SDC supplemented with either 2,5- or 1,6-hexanediol (5% final concentration), and cells were imaged at 39**°**C for the times indicated.

The images were acquired across 8 to 10 planes on the z-axis with an interplanar distance of 0.25 μm. The dual-color images were captured in the sequential scanning mode for minimizing crosstalk between fluorophore channels. Post-acquisition, images were deconvolved by YacuDecu function that utilizes Richardson-Lucy algorithm for calculating theoretical Point Spread Functions (PSFs) (https://github.com/bnorthan) (Rueden et al., 2017). The PSFs were computed each time based on the set of microscopy parameters used in the imaging analyses. Custom plugins for Fiji (Schindelin et al., 2012) were used to colorize, split or merge channels, make composites and adjust brightness of the images. 3D rendering and visualization were performed using either ClearVolume (Royer et al., 2015) or arivis Vision 4D software v. 3.1 (render mode: maximum intensity; arivis AG, Rostok, Germany).

For fixed-cell imaging, cells were grown at 30°C to early log phase in YPD, subjected to instantaneous heat shock at 39°C for the indicated times, and then fixed in 1% formaldehyde for 7 min. Cells were harvested and the pellet was washed with 1X phosphate-buffered saline (PBS) (pH 7.4). An aliquot of cells was immobilized onto concanavalin A-coated glass bottom dish, and then imaged using Leica TCS SP5 II STED-CW laser scanning confocal microscope (Leica Microsystems, Inc., Buffalo Grove, IL). Images were acquired and processed as above.

For colocalization analyses, the fractional overlap metric scores (Manders’ colocalization coefficients) were calculated using JACoP plugin implemented in ImageJ (Bolte and Cordelieres, 2006; Schneider et al., 2012). The average intensity plots were generated using the plot profile feature in ImageJ. Intensities of pixels were obtained along the line path (as indicated in Figure 1G) for each z-section. Intensities across nine individual z-sections were combined and the average was plotted for each channel.

For analysis of gene coalescence, nine z-planes with an interplanar distance of 0.5 μm, covering the entire depth of the nucleus, were inspected for location of tagged genomic loci. The relative nuclear positions of the tagged loci were assessed by measuring 3D distances between them. The 3D distances were measured as 3D polyline lengths between the signal centroids (green and red spots). A cell was scored positive for coalescence if the 3D distance between the centroids of green and red spots was between 0.3 and 0.7 μm. 3D distance measurements, 3D rendering and visualization were performed in arivis Vision4D.

### Super-resolution microscopy

Yeast cells were grown at 30°C to early log phase (A_600_ = 0.5) in synthetic dextrose complete (SDC) medium. An aliquot of living cells was immobilized onto concanavalin A-coated glass bottom dish. Fresh SDC medium was added in the dish before imaging. Samples were subjected to heat shock (39**°**C) by heating the objective using Bioptechs objective heater system and by controlling the temperature of incubator chamber enclosing the microscope. High-resolution Stimulated Emission Depletion (STED) images were acquired on Leica TCS SP5 II STED-CW laser scanning confocal microscope (Leica Microsystems, Inc., Buffalo Grove, IL). Emission depletion was accomplished with 592 nm STED laser. Images were deconvolved as above, rendered (render mode: volumetric) and visualized in arivis Vision4D.

### Hsf1, Med15 and Rpb1 ChIP-seq data analysis

Hsf1 ChIP-seq data were obtained from GSE117653 (Pincus et al., 2018). Med15 data are from PRJNA657372 (Sarkar et al., 2020). Rpb1 ChIP-seq data were obtained from GSE125226 (Albert et al., 2019). Reads were aligned to the S. cerevisiae reference genome (SacCer3) using Bowtie 2 (Langmead and Salzberg, 2012). SAM files were converted to BAM format using SAMtools (Li et al., 2009). BAM files were then converted to bigWig and bedGraph format at bin size of 1 (-bs 1) and normalized to the library size (--normalizeUsing CPM) using bamCoverage function of deepTools2 (Ramirez et al., 2016). The bigWig files were used to obtain genome browser tracks in Integrative Genomics Viewer (IGV) browser (Robinson et al., 2011), and to make metagene plots. Metagene profiles were created using computeMatrix and plotProfile function tools in deepTools2. The plots are scaled +/-1 kb of ORFs of the Hsf1-dependent genes (Pincus et al., 2018). Occupancy of Hsf1, Med15 and Rpb1 (Pol II) were computed using the bedtools map function. The occupancy of Hsf1 and Med15 were obtained as sum of signals across 1 kb region upstream of ORFs of protein-coding genes. For occupancy of Pol II, sum of signals across ORFs were normalized to length of the ORFs.

### Nascent transcript sequencing data analysis

NAC-seq data were obtained from GSE117653 (Pincus et al., 2018). Reads were aligned to the S. cerevisiae reference genome (SacCer3) using TopHat2 (Kim et al., 2013) with --segment-length 20 -I 2500 options. Wiggle files were generated by normalizing to the library size (--normalizeUsing CPM) using the bamCoverage function of deepTools2 and visualized in IGV genome browser.

### Analysis of average relative localization of factors

Foci of LacO-tagged HSP104 gene locus or MS2-tagged HSP12 mRNA locus were manually identified in a specific z plane and centered within a square of 12 x 12 pixels. Distribution of signal from other channel, depicting distribution of secondary factor, was gathered from the corresponding 12 x 12 squares in the same z plane. A composite text image was created by computing average of centered images from roughly 30 cells per condition. The text images were used to generate contour plots. The contour plots were created using the Plotly function (Plotly Technologies Inc., 2015) in R (R Core Team, 2020). The intensity minima and maxima, spanning the entire range of signals in the given channel, were split into 800 steps.

### Quantification of 3C

TaqI-3C data was quantified as described in (Chowdhary et al., 2020; Chowdhary et al., 2019). The percent digestion efficiency was determined by amplifying a region across Taq I restriction site using a pair of convergent primers (sequences provided in Table S3). The percent digestion efficiencies were determined for each primer combination and incorporated into the following formula:

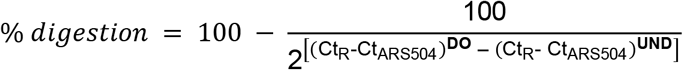

Here, Ct_R_ is the cycle threshold (Ct) quantification of the digested only (DO) or undigested (UND) templates, and Ct_ARS504_ the cycle threshold quantification of the ARS504 locus (a region lacking a Taq I site).

For measurement of normalized frequency of intragenic or intergenic interactions, the Ct values for digested only (DO_3C_) and ligated (Lig_3C_) templates for crosslinked chromatin, and digested only and ligated genomic DNA (Lig_gDNA_ and DO_gDNA_, respectively) were incorporated into the following formula:

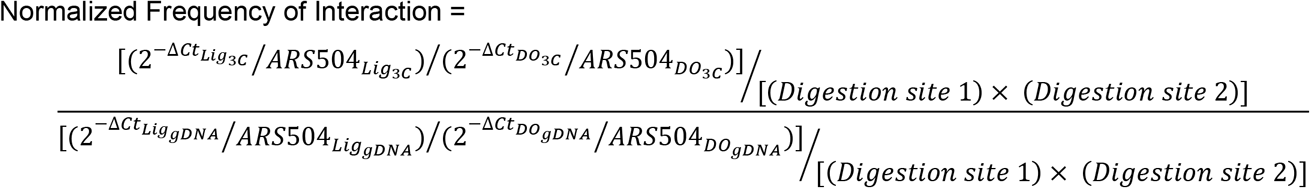

i. ΔCt values were obtained by subtracting Ct (no-template) from those of Lig_3C_, DO_3C_, or gDNA templates.
ii. 2^-ΔCt^?RS504 are the fold-over signals normalized to *ARS504* locus.
iii. Ligation-dependent signals (LDS) are determined as ratio of fold-over normalized signals of Lig_3C_ and DO_3C_ templates; also applicable to the gDNA control.

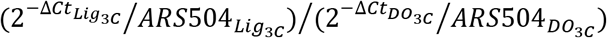
iv. LDS were corrected for variation in Taq I digestion efficiencies of sites 1 and 2 (as detailed above).
v. Normalized Frequency of Interaction is defined as the ratio of ligation-dependent signals of 3C and gDNA control templates after correcting for differences in their digestion efficiencies.

### Statistical tests used

Student’s t test (two-tailed) was used to calculate statistical significance between all pairwise comparisons (as assumptions of parametric distributions were fulfilled), except in Figure 2C, one-way ANOVA followed by Tukey’s post hoc analysis was used. Each pairwise comparison is done with means of two independent biological samples (N=2) +SD. n.s., P>0.05; *, P<0.05; **, P<0.01; ***, P<0.001.

## AUTHOR CONTRIBUTIONS

Conceptualization: S. C., D. S. G. and D. P.; Methodology: S. C., A. S. K., D. S. G. and D. P; Formal Analysis: S. C. and A. S. K.; Investigation: S. C. and A. S. K.; Resources: S. C., A. S. K., S. P.; Writing – original draft: S. C. and D. P.; Writing – Review and Editing: S. C., A. S. K., S. P., D. S. G. and D. P.; Visualization: S. C., A. S. K. and D. P.; Supervision: D. S. G. and D. P.; Funding Acquisition: D. S. G. and D. P.

## ACKNOWLEDGEMENTS

We would like to thank Vytas Bindokas and Christine Labno at the University of Chicago Integrated Light Microscopy Core (RRID: SCR_019197) for imaging assistance and technical support. We thank Rama Rangana than and Alex Ruthenburg for sharing reagents and equipment. We also thank Luke Dyer for help with strain construction and Pincus and Gross lab members for helpful discussions. This work was supported by NIH grants R01 GM138689 to D.P. and R01 GM138988 and R15 GM128065 to D.S.G.

