## Supplemental information for "Inducible transcriptional condensates drive 3D genome reorganization in the heat shock response"

### SUPPLEMENTAL INFORMATION
**Chowdhary et al.**

Supplemental Figures (S1-S4)

Supplemental Tables (S1-S6)

### Figure S1

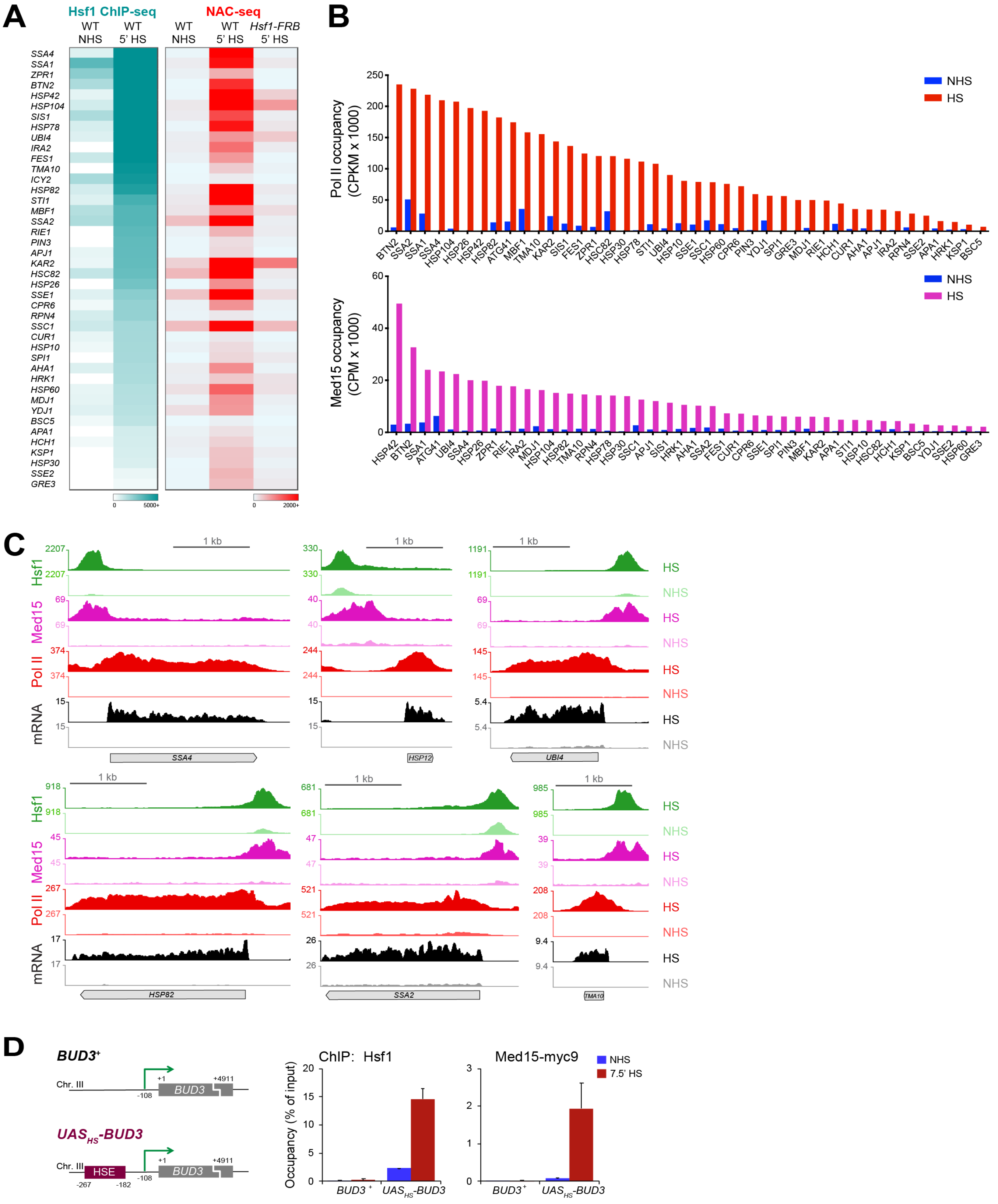

A
Hsf1 ChIP-seq
NAC-seq
WT NHS
WT 5' HS
WT NHS
WT 5' HS
Hsf1-FRB 5' HS
SSA4
SSA1
ZPR1
BTN2
HSP42
HSP104
SIS1
HSP78
UBI4
IRA2
FES1
TMA10
ICY2
HSP82
STI1
MBF1
SSA2
RIE1
PIN3
APJ1
KAR2
HSC82
HSP26
SSE1
CPR6
RPN4
SSC1
CUR1
HSP10
SPI1
AHA1
HRK1
HSP60
MDJ1
YDJ1
BSC5
APA1
HCH1
KSP1
HSP30
SSE2
GRE3
0
5000+
0
2000+
B
Pol II occupancy (CPKM x 1000)
NHS
HS
Med15 occupancy (CPM x 1000)
C
Hsf1
Med15
Pol II
mRNA
1 kb
HS
NHS
SSA4
HSP12
UBI4
HSP82
SSA2
TMA10
D
BUD3+
Chr. III
+1
+4911
-108
BUD3
UAS_HS-BUD3
HSE
-267
-182
ChIP: Hsf1
Med15-myc9
NHS
7.5' HS
Occupancy (% of input)
BUD3+
UAS_HS-BUD3

**Figure S1. Heat shock genes associate with exceptionally high levels of Hsf1, Mediator and RNA Pol II**

**A)** Heat Shock Factor 1 (Hsf1) occupancy and nascent-RNA transcription at genes bound and induced by Hsf1 in S. cerevisiae (n=42). Left: heatmaps depicting occupancy of Heat Shock Factor 1 (Hsf1) in non-heat shock (NHS) and 5 min heat shock (HS) conditions, as determined by ChIP-seq (gray to teal). Right: heatmaps showing nascent-RNA densities in presence and absence of heat shock and Hsf1, as determined by NAC-seq (blue to red). Genes are sorted in the descending order of Hsf1 ChIP-seq counts (top to bottom). NAC-seq and Hsf1 ChIP-seq data were derived from (Pincus et al., 2018).
**B)** RNA Pol II (top) and Med15 (bottom) occupancy at Hsf1-dependent genes in NHS and HS conditions. For Med15 ChIP-seq analysis, cells were heat-shocked for 15 min at 37ºC (Sarkar et al., 2019). For Pol II, 5 min at 40ºC (Albert et al., 2019).
**C)** IGV browser views of Hsf1 (green), Med15 (magenta) and RNA Pol II (Rpb1; red) ChIP-seq densities as well as NAC-seq counts (black) at the indicated genes in NHS and HS conditions.
**D)** Left: physical maps of *BUD3* and the chromosomal transgene $UAS_{HS}$-*BUD3*. Right: Hsf1 and Med15-myc9 occupancy at $BUD3^{+}$ and $UAS_{HS}$-*BUD3* in NHS and 7.5 min HS conditions (n=2; qPCR=4).

### Figure S2

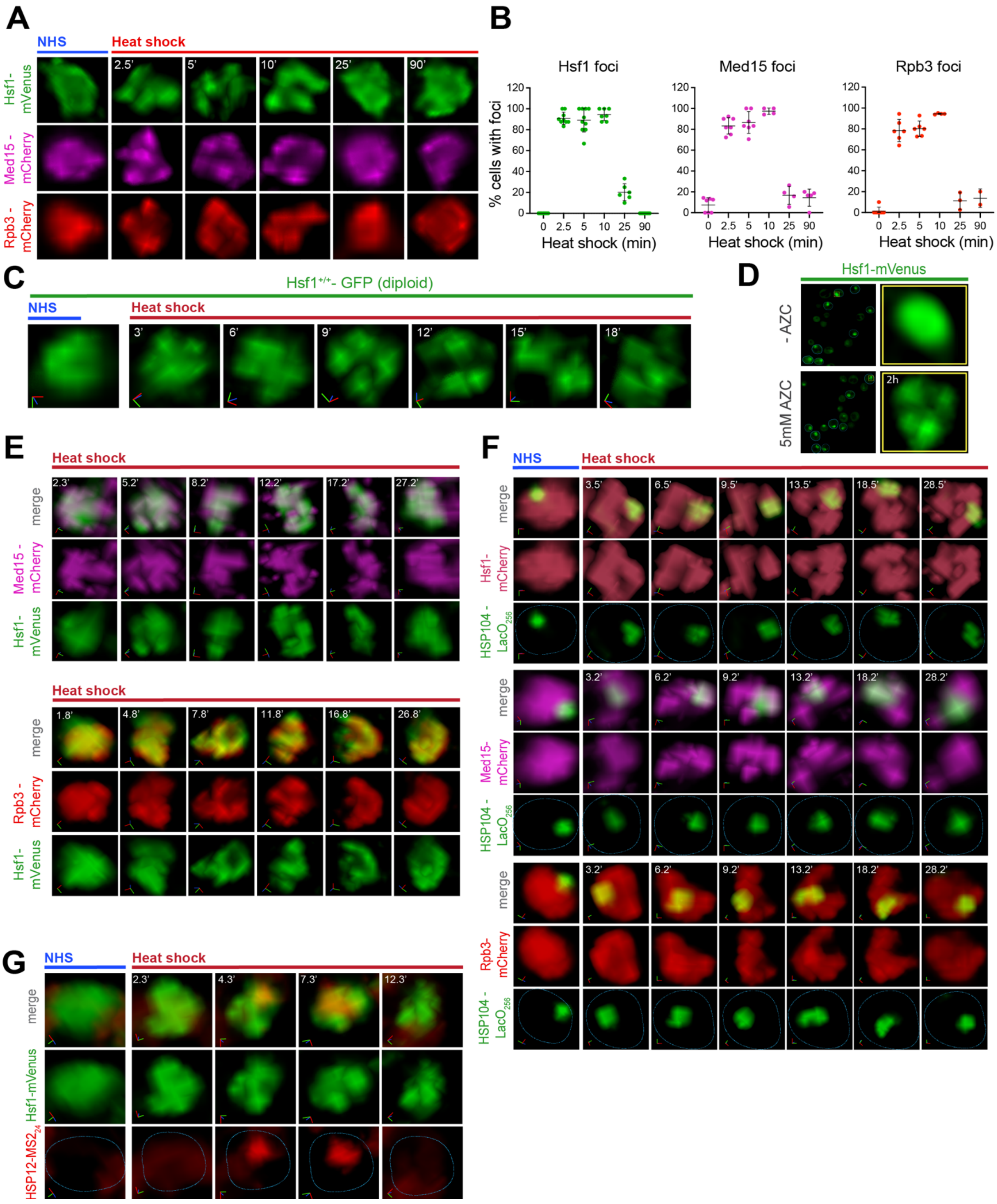

A
NHS
Heat shock
2.5'
5'
10'
25'
90'
Hsf1-mVenus
Med15 - mCherry
Rpb3 - mCherry
B
Hsf1 foci
Med15 foci
Rpb3 foci
% cells with foci
Heat shock (min)
C
Hsf1+/+- GFP (diploid)
3'
6'
9'
12'
15'
18'
D
Hsf1-mVenus
- AZC
5mM AZC
2h
E
merge
F
HSP104 - LacO256
G
HSP12-MS2

**Figure S2. Hsf1 forms transcriptionally active clusters upon stress**

**A)** Fixed-cell microscopy of cells expressing Hsf1-mVenus (top), Med15-mCherry (middle) or Rpb3-mCherry (bottom) in presence or absence of heat shock. Cells were fixed with formaldehyde before or following HS, then imaged. Shown are enlarged nuclei of representative cells at each time point and condition. 3D volumes were rendered and visualized in ClearVolume. For Med15-mCherry, z=7 (25' and 90' HS).
**B)** Percentage of cells expressing Hsf1 (left), Med15 (middle) or Rpb3 (right) foci. Left: cells expressing Hsf1-mVenus were fixed with formaldehyde before or following HS and imaged. Cells with >2 foci were counted as those containing foci. A total of 50-90 cells were evaluated per time point. Shown are means +/- SD. Middle and right: same as left, except shown for cells expressing Med15-mCherry or Rpb3-mCherry. 80-90 cells were evaluated per time point for each strain type.
**C)** Live-cell imaging of diploid strain ASK741 expressing Hsf1-GFP before (NHS, 26.5ºC) and following heat shock (39ºC) for the times (t) indicated. A single cell is followed throughout the heat shock time course. Shown is the 3D volumetric rendering of each nucleus; x (red), y (green) and z (blue) axes are indicated. For HS images, z=11.
**D)** Live imaging of cells expressing Hsf1-mVenus in presence or absence of L-azetidine-2-carboxylic acid (AZC). Left: representative images of cells expressing Hsf1-mVenus before or following 2h of AZC treatment. Blue line highlights cell boundary. Right: enlarged view of the nucleus indicated in yellow boxes on the left. Cells were rendered and visualized by ClearVolume.
**E)** Live imaging of cells co-expressing Hsf1-mVenus and Med15-mCherry (top), or Hsf1-mVenus and Rpb3-mCherry (bottom). Shown are enlarged 3D rendered nuclei of representative cells that were imaged following heat shock for indicated times. Note that in each case, a single cell is followed through the heat shock time course. Images were rendered and visualized in arivis.
**F)** 3D rendered micrographs of Hsf1-mCherry (top), Med15-mCherry (middle) or Rpb3-mCherry (bottom) and the *HSP104-LacO*$_{256}$ gene locus before and following heat shock for the indicated times. A single cell is followed through the heat shock time course. Blue dotted line highlights nuclear boundary. 3D rendering and visualization was performed in arivis; x (red), y (green) and z (blue) axes are shown.
**G)** 3D rendered micrographs of Hsf1-mVenus and the MCP-mCherry labeled *HSP12-MS2*$_{24}$ mRNA before or following heat shock for the indicated times. Blue dotted line highlights nuclear boundary. A single cell is followed throughout the heat shock time course. 3D rendering and visualization was performed in arivis.

### Figure S3

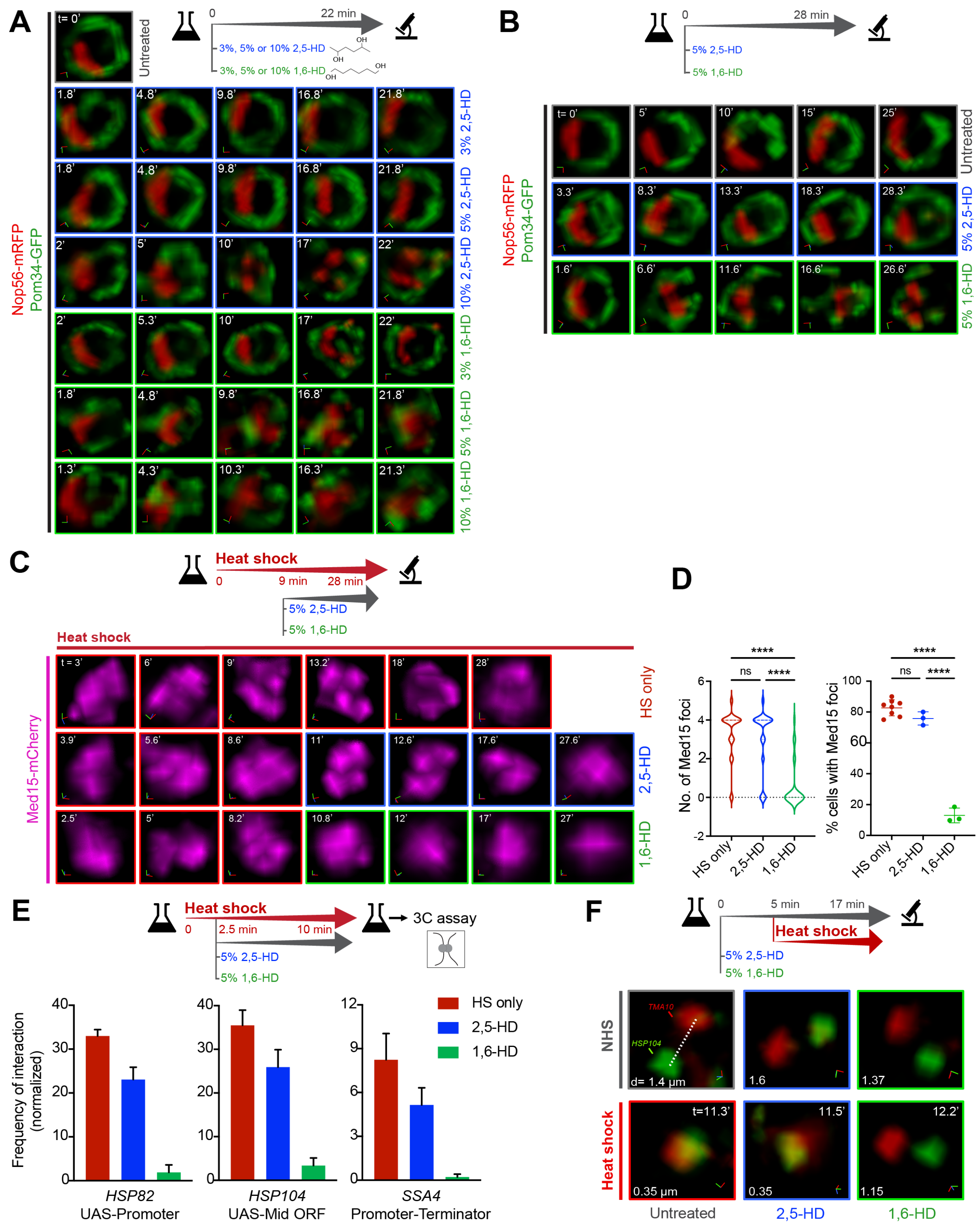

A
t= 0'
Untreated
0
22 min
3%, 5% or 10% 2,5-HD
3%, 5% or 10% 1,6-HD
OH
Nop56-mRFP
Pom34-GFP
1.8'
4.8'
9.8'
16.8'
21.8'
3% 2,5-HD
5% 2,5-HD
2'
5'
10'
17'
22'
10% 2,5-HD
5.3'
3% 1,6-HD
5% 1,6-HD
1.3'
4.3'
10.3'
16.3'
21.3'
10% 1,6-HD
B
0
28 min
5% 2,5-HD
5% 1,6-HD
t= 0'
25'
15'
Untreated
3.3'
8.3'
13.3'
18.3'
28.3'
1.6'
6.6'
11.6'
16.6'
26.6'
C
Heat shock
0
9 min
28 min
5% 2,5-HD
5% 1,6-HD
Med15-mCherry
t = 3'
6'
9'
13.2'
18'
28'
HS only
3.9'
5.6'
8.6'
11'
12.6'
17.6'
27.6'
2,5-HD
2.5'
8.2'
10.8'
12'
27'
1,6-HD
D
No. of Med15 foci
% cells with Med15 foci
****
ns
HS only
E
2.5 min
10 min
3C assay
Frequency of interaction (normalized)
HSP82
UAS-Promoter
HSP104
UAS-Mid ORF
SSA4
Promoter-Terminator
F
5 min
17 min
NHS
TMA10
HSP104
d= 1.4 µm
1.6
1.37
t=11.3'
11.5'
12.2'
0.35 µm
0.35
1.15
Untreated

**Figure S3. 1,6-hexanediol disrupts the Nucleolus, the nuclear pore complex, Mediator clusters and gene interactions within and between Hsf1 target genes**

**A)** 3D rendered micrographs of representative cells in absence of alcohol (untreated) or treated with either 2,5-HD or 1,6-HD at 3, 5 or 10% concentrations. Row 1, right: experimental workflow. Rows 1-7: Live-cell imaging of diploid strain SCY712 (Nop56-RFP Pom34-GFP). Red, Nop56-RFP (component of the nucleolus); Green, Pom34 (component of the nuclear pore complex). z =6. Note that in each row, a single cell is followed throughout the treatment.
**B)** Comparison of effects of 5% 2,5-HD vs. 5% 1,6-HD on the nucleolus and nuclear pore complex. Analysis and presentation as in A; z =7.
**C)** 3D rendered micrographs of a representative cell subjected to heat shock for times (t) indicated in the absence of alcohol (top) or treated with either 2,5-HD (middle) or 1,6-HD (bottom) after 9 min of HS. Note that a different cell was imaged after alcohol treatment.
**D)** Number of Med15 foci (left) and the percentage of cells showing >2 foci (right) in presence or absence of HD. Cells expressing Med15-mCherry were heat-shocked for 9 min followed by treatment with either 2,5- or 1,6-hexanediol (or not). Cells were imaged after ~3 min of adding the drug. 100-120 cells were evaluated per condition. ****P <0.0001; ns (not significant), P >0.05. P values were calculated by ANOVA followed by Tukey's post hoc analysis.
**E)** Intragenic interactions within indicated Hsf1 target genes as determined by TaqI-3C assay. Top: experimental workflow. Bottom: Intragenic contact frequencies detected within *HSP82, HSP104* and *SSA4* in cells heat-shocked and treated with either 2,5- or 1,6-hexanediol (or not). Depicted are means +SD (n=2; qPCR=4).
**F)** Intergenic interactions between *lacO*-tagged *HSP104* and *tetO*-tagged *TMA10* gene loci as determined by live cell imaging analysis. Top: experimental workflow. Cells were pre-treated with either 5% 2,5- or 1,6-hexanediol (or not) for 5 min followed by heat shock of 12 min (or not) and imaged at times (t) indicated. Middle: representative images of cells that were pre-treated with either 1,6-HD or 2,5-HD or left untreated (right to left). Bottom: same as above, except cells were subjected to heat shock following treatment with alcohol (or not). Cells were rendered and visualized in arivis; x (red), y (green) and z (blue) axes are shown. d, 3D distances measured between signal centroids. Images were taken across 9 planes in z-direction; step size =0.5 µm.

### Figure S4

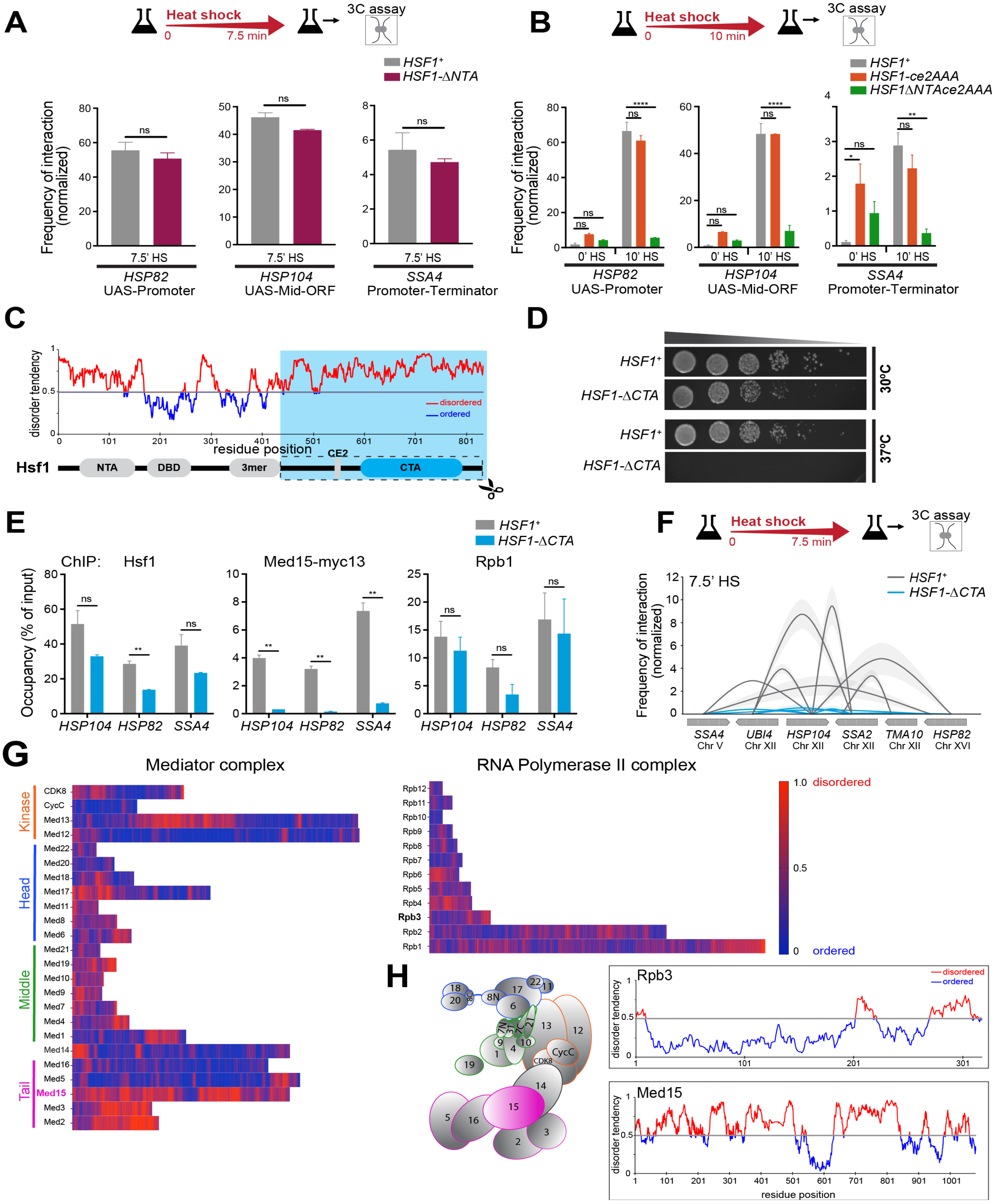
A
Heat shock
0
7.5 min
3C assay
HSF1+
HSF1-ΔNTA
Frequency of interaction (normalized)
7.5' HS
HSP82
UAS-Promoter
HSP104
UAS-Mid-ORF
SSA4
Promoter-Terminator
ns
B
Heat shock
0
10 min
3C assay
HSF1+
HSF1-ce2AAA
HSF1ΔNTAce2AAA
0' HS
10' HS
C
disorder tendency
residue position
disordered
ordered
Hsf1
NTA
DBD
3mer
CE2
CTA
D
HSF1+
HSF1-ΔCTA
30°C
37°C
E
ChIP: Hsf1
Med15-myc13
Rpb1
Occupancy (% of input)
HSP104
HSP82
SSA4
F
Heat shock
0
7.5 min
3C assay
7.5' HS
SSA4 Chr V
UBI4 Chr XII
HSP104 Chr XII
SSA2 Chr XII
TMA10 Chr XII
HSP82 Chr XVI
G
Mediator complex
RNA Polymerase II complex
Kinase
Head
Middle
Tail
CDK8
CycC
Med13
Med12
Med22
Med20
Med18
Med17
Med11
Med8
Med6
Med21
Med19
Med10
Med9
Med7
Med4
Med1
Med14
Med16
Med5
Med15
Med3
Med2
Rpb12
Rpb11
Rpb10
Rpb9
Rpb8
Rpb7
Rpb6
Rpb5
Rpb4
Rpb3
Rpb2
Rpb1
1.0 disordered
0.5
0 ordered
H
Rpb3
Med15
disorder tendency
residue position
disordered
ordered

**Figure S4. Stable Mediator occupancy is necessary for driving intergenic interactions between Hsf1-target genes**

**A)** Intragenic looping and crumpling interactions within indicated Hsf1 target genes in WT and *HSF1-ΔNTA* cells heat-shocked for 7.5 min. Top: experimental workflow of TaqI-3C assay. Bottom: Intragenic contact frequencies detected within *HSP82, HSP104* and *SSA4* in 7.5 min-HS conditions. Depicted are means +SD (n=2; qPCR=4). ns (not significant), P >0.05 (calculated using two-tail t test)
**B)** Intragenic interactions within indicated Hsf1 target genes in *WT, HSF1-ce2AAA* and *HSF1-ΔNTAce2AAA* cells under NHS and 10 min-HS conditions. Top: experimental workflow of TaqI-3C assay. Bottom: Intragenic contact frequencies detected within *HSP82, HSP104* and *SSA4* in NHS and 10min-HS conditions. Depicted are means +SD (n=2; qPCR=4) ****P <0.0001; **P <0.01; *P <0.05; ns (not significant), P >0.05. P values were calculated by ANOVA followed by Tukey's post hoc analysis.
**C)** Domain map and prediction of disorder in Hsf1. Top: predicted value of disorder tendency for each amino acid within Hsf1 obtained using IUPRED. Values >0.5 (red) are disordered and values <0.5 (blue) are ordered. Bottom: Domain map of Hsf1. NTA, N-terminal activation domain; DBD, DNA binding domain; 3mer, trimerization domain; CE2, conserved element 2 domain; CTA (blue), C-terminal activation domain. Box: truncation mutation in the Hsf1-ΔCTA mutant.
**D)** Spot dilution analysis of *HSF1*$^{+}$ and *HSF1-ΔCTA* cells. Five-fold serial dilutions of cells were spotted onto YPD. Plates were incubated at 30ºC or 37ºC for 30 h.
**E)** Hsf1, Med15-myc13 and Rpb1 ChIP analysis of *HSP104, HSP82* and *SSA4*. Wild type and *Hsf1-ΔCTA* strains were heat-shocked for 7.5 min, then processed for ChIP. Depicted are means + SD (n=2; qPCR=4). **, P <0.01; ns (not significant), P >0.05 (calculated using two-tail t-test).
**F)** Intergenic contacts (solid arcs) between indicated Hsf1 target gene pairs for *HSF1*$^{+}$ and *HSF1-ΔNTA* strains. Cells were heat-shocked for 7.5 min, then processed for TaqI-3C. Values indicate normalized interaction frequencies. Gene regions are depicted as UAS, promoter, mid-ORF and terminator (from blunt to arrowhead direction). Data are derived from two independent biological replicates; qPCR=4. Depicted are means +/- SD (shaded region around solid arcs).
**G)** *In silico* analysis of disorder prediction in the Mediator and RNA Pol II subunits. Shown are heat maps of predicted values of disorder for each amino acid of a subunit. Heat maps are presented from amino to carboxyl terminus (left to right). Disorder tendencies were calculated using IUPRED. Values <0.5 (blue) are ordered and >0.5 (red) are disordered.
**H)** Left: yeast Mediator subunits in Head (blue), Middle (green), Tail (magenta) and Kinase (orange) modules. Med14 is a scaffold. Right: prediction of disorder in Med15 and Rpb3 protein subunits of the Mediator and RNA Pol II complexes, respectively. The plots score predicted values of disorder tendency for each amino acid within Med15 or Rpb3, as per IUPRED. Values <0.5 (blue) are ordered and >0.5 are disordered (red).

**Table S1. Yeast strains**

| Strain name | Genotype | Reference/ Source |
|---|---|---|
| DPY001 | *MATa ADE2 trp1-1 can1-100 leu2-3,112 his3-11,15 ura3-1* | El-Samad lab |
| DPY032 | DPY001; *HSF1-mVenus::HIS3* | This study |
| SCY001 | DPY001; *MED15-mCherry::hphMX6* | This study |
| SCY002 | DPY001; *RPB3-mCherry::hphMX6* | This study |
| SCY003 | DPY032; *MED15-mCherry::hphMX6* | This study |
| SCY004 | DPY032; *RPB3-mCherry::hphMX6* | This study |
| DBY255 | *MATa ade2-1 can1-100 leu2-3,112 trp1-1 ura3-1* his3-11,15::GFP-LacI::HIS3 *HSP104-LacO$_{256}$::TRP1 SEC63-MYC*×13*::KAN-MX* | Brickner et al., 2012 |
| SCY005 | DBY255; *HSF1-mCherry::hphMX6* | This study |
| SCY006 | DBY255; *MED15-mCherry::hphMX6* | This study |
| SCY007 | DBY255; *RPB3-mCherry::hphMX6* | This study |
| SCY008 | DPY032; *ura3-1::MCP-mCherry::URA3* | This study |
| SCY009 | SCY008; *HSP12-MS2$_{24}$-loxp-KAN-MX6-loxp* | This study |
| SCY010 | SCY009; *pSH69-Cre-hphMX6* | This study |
| SCY011 | SCY010*; HSP12-MS2$_{24}$-loxp* | This study |
| BY4741 | *MATa his3Δ1 leu2Δ0 met15Δ0 ura3Δ0* | Chowdhary et al., 2019 |
| DSG144 | BY4741; *trp1Δ::KAN-MX* | Gross Lab |
| ASK804 | BY4741; *3xHSE-BUD3* | Chowdhary et al., 2019 |
| LRY003 | ASK804*; trp1Δ::KAN-MX* | This study |
| ASK213 | DSG144; *MED15-MYCx9::TRP1* | This study |
| ASK214 | LRY003; *MED15-MYCx9::TRP1* | This study |
| ASK741 | MATα/MATa; Hsf1-GFP::HIS3MX6/ Hsf1-GFP::HIS3MX6 | This study |
| ASK727 | *MAT*α/MATa *ade2-1/ ade2-1 CAN1$^{+}$/can1-100 ura3-1/ ura3-1* THR$^{+}$/thr1-4 *leu2-3,112/ leu2-3,112 trp1-1/ trp1$^{-}$* his3-11,15::GFP-LacI::HIS3/ his$^{-}$ *HSP104-LacO$_{256}$::TRP1/HSP104$^{+}$ TMA10-tetO$_{200}$::LEU2/ TMA10$^{+}$ leu2::tetR-mCherry::hphMX::leu2 SEC63-MYC*×13 | Chowdhary et al., 2019 |
| SCY712 | *MATa/*MATα his3Δ1/his3Δ1 leu2Δ0/leu2Δ0 l*ys2Δ0/ LYS$^{+}$* met15*Δ*0/MET$^{+}$ *ura3Δ0/ ura3Δ0 SIK1-mRFP::KanMX6/SIK1$^{+}$* POM34-GFP::HIS3MX6/POM34$^{+}$ | Chowdhary et al., 2019 |

| | | |
|---|---|---|
| DPY182 | DPY001; *hsf1∆::KAN; HSF1pr-HSF1-GFP::TRP1* | This study |
| DPY034 | DPY001; *hsf1∆::KAN; pRS316-HSF1* | This study |
| DPY1805 | DPY001; *trp1-1::HSF1pr-HSF1-ce2AAA-GFP::TRP1* | This study |
| SCY012 | DPY1805; *MED15-mCherry::hphMX6* | This study |
| SCY013 | DPY1805; *RPB3-mCherry::hphMX6* | This study |
| DPY144 | DPY001*; HSE-YFP::LEU2* | Krakowiak et al., 2018 |
| ASK215 | DPY144; *MED15-MYCx13::HIS3* | This study |
| DPY304 | DPY001*; HSE-YFP::LEU2 hsf1∆::KAN HSF1pr-HS-F1::TRP1* | Krakowiak et al., 2018 |
| DPY417 | DPY001*; HSE-YFP::LEU2 hsf1∆::KAN HSF1pr-hs-f1(147-833)::TRP1* | Krakowiak et al., 2018 |
| ASK216 | DPY417; *MED15-MYCx13::HIS3* | This study |
| DPY179 | DPY001; *hsf1∆::KAN HSF1pr-hsf1(147-833)-GF-P::TRP1* | This study |
| SCY014 | DPY179; *MED15-mCherry::hphMX6* | This study |
| SCY015 | DPY179; *RPB3-mCherry::hphMX6* | This study |
| DPY1818 | DPY001; *trp1-1::HSF1pr-hsf1(147-833)-ce2AAA-GFP::TRP1* | This study |
| SCY016 | DPY1818; *MED15-mCherry::hphMX6* | This study |
| SCY017 | DPY1818; *RPB3-mCherry::hphMX6* | This study |
| DPY418 | DPY001*; HSE-YFP::LEU2 hsf1∆::KAN HSF1pr-hs-f1(1-424)::TRP1* | Krakowiak et al., 2018 |
| ASK217 | DPY418; *MED15-MYCx13::HIS3* | This study |

**Table S2. Forward (F) primers used in 3C analysis**

| Name | Sequence (5’ → 3’) |
|---|---|
| UBI4 F+524 | GTAAGCAGCTAGAAGATGGTAGAACC |
| UBI4 F+1624 | TGATACGGATAGAATATTGTGACTACC |
| HSP104 F-63 | AGGCATTGTAATCTTGCCTCAATTC |
| HSP104 F+782 | GTAAGACCGCTATTATTGAAGGTG |
| HSP104 F+1550 | CCCTTGATGCTGAACGTAGATATG |
| HSP104 F+2756 | AGGTGATGACGATAATGAGGACAG |
| SSA2 F-242 | CACTGCATTCTTACTCTCTCTTGG |
| SSA2 F+198 | AGGTAACAGAACCACTCCATCTTTC |
| SSA2 F+1368 | TCTCTACTTATGCTGACAACCAACC |
| SSA2 F+2297 | AGTGACTTGAAGACTAGGAATATCG |
| TMA10 F+159 | ACGAAGCAAAGTCTAACCCAAAG |
| TMA10 F+811 | ATGCAAAAACACTTCCCAGAATAG |
| SSA4 F-268 | ACACGAAAGATATCTCAACTCTAGCC |

| | |
|---|---|
| SSA4 F+198 | GCCTTCTTATGTGGCTTTTACTGAC |
| SSA4 F+2255 | ATAAGAAAGTCATCGCCAAACAAC |
| HSP12 F-47 | ACGTATAAATAGGACGGTGAATTGC |
| HSP12 F+279 | AAAAGGCAAGGATAACGCTGAAG |
| HSP82 F-290 | CCTCTCTCAACACAGTAATCCATAAAC |
| HSP82 F+740 | AATTAGTCGTCACCAAGGAAGTTG |
| HSP82 F+1445 | GCCAGAACACCAAAAGAACATCTAC |
| HSP82 F+2189 | ATGAGGATGAAGAAACAGAGACTGC |
| ARS504F (internal control) | GTCAGACCTGTTCCTTTAAGAGG |

**Table S3. Reverse (R) primer used for percent digestion determination in 3C**

| Name | Sequence (5' → 3') |
|---|---|
| UBI4 R+524 | TGAATTTTCGACTTAACGTTGTCG |
| UBI4 R+1624 | ATCACCCAGTATCCCTGATTTCAC |
| HSP104 R-63 | ATCGTTAGAGCCCTTTCTGTAAATTG |
| HSP104 R+782 | TTCTTCGATTTCCTTCAAAACACC |
| HSP104 R+1550 | CCACATTTTGGATCATGGAGTTG |
| HSP104 R+2756 | TCTTTTGCTCGGGTGTCAAGTTC |
| SSA2 R-242 | GATGGAATGTTCTAGAAAAAAACTTC |
| SSA2 R+198 | GCTTCATATCACCTTGGACTTCTG |
| SSA2 R+1368 | TTCAATTTGTGGGACACCTCTTG |
| SSA2 R+2297 | GACGCCCTTACGAATAGAACTTTAC |
| TMA10 R+159 | TTTCATTGTTTTGGGAGTTAGAGC |
| TMA10 R+811 | CCGGTTATAGGACCCTTATTGATG |
| SSA4 R-268 | TGTTACTGTCGTCAAACTAAGGAG |
| SSA4 R+198 | TTTACGTCCGATCAGACGCTTAG |
| SSA4 R+2255 | GTGTTAAACTCCGGTCAAAAGAAAC |
| HSP12 R-47 | TTCAGAAGCTTTTTCACCGAATC |
| HSP12 R+279 | AAACCATGTAACTACAAAGAGTTCCG |
| HSP82 R-290 | GAAGGACCTGGTTGGTATTAAGATG |
| HSP82 R+740 | AATGCTTAACGTACAATGGGTCTTC |
| HSP82 R+1445 | ATTCATCAATTGGGTCGGTCAAG |
| HSP82 R+2189 | ACACACTAGACGCGTCGGAATAG |
| ARS504R (internal control) | CATACCCTCGGGTCAAACAC |

**Table S4. Primers used in ChIP analysis**

| Name | Sequence (5' ® 3') |
|---|---|
| BUD3 UAS F (-294 to -72) | GCTCTTTGTCATACGCATAGAATTG |
| BUD3 UAS R | CAGTAGAATGCGAGTACAGACAAAC |
| HSP104 UAS F (-266 to -195) | CTTAAACGTTCCATAAGGGGC |
| HSP104 UAS R | TGCAGTTCTTTGAGATGGGCC |
| HSP104 PROM F (-132 to +41) | AGGCATTGTAATCTTGCCTCATTC |
| HSP104 PROM R | ATCGTTAGAGCCCTTTCTGTAAATTG |
| HSP82 UAS F (-393 to -155) | CCTCTCTCAACACAGTAATCCATAAAC |
| HSP82 UAS R | GAAGGACCTGGTTGGTATTAAGATG |
| HSP82 PROM F (-156 to -88) | TCCGCCACCCCCTAAAAC |
| HSP82 PROM R | TGAGGAGGTCACAGATGTTAA |
| SSA4 UAS F (-396 to -145) | ACACGAAAGATATCTCAACTCTAGCC |
| SSA4 UAS R | TGTTACTGTCGTCAAACTAAGGAG |
| SSA4 PROM F (-245 to +35) | AGTTCCTAGAACCTTATGGAAGCAC |
| SSA4 PROM R | GTTGTACCTAAATCAATACCAACAGC |

**Table S5. Primer used in RT-qPCR analysis**

| Name | Sequence (5' → 3') |
|---|---|
| HSP104 F+1922 | TTAGCTAATCCAAGGCAACCAG |
| HSP104 ORF R+1922 | ACCTTCATCGTACCCGACATAAC |
| HSP82 F+1838 | ACATGGAAAGAATCATGAAGGCTC |
| HSP82 R +1838 | AAGTCCTTGACAGTCTTGTCTTGAG |
| SSA4 F+1539 | ATCTACTGGGTAAATTTGAGTTGAGC |
| SSA4 R+1539 | CTAGCTGATTCTTAGCTTGAACACG |
| SSA2 F+1905 | CTAAATTGTACCAAGCTGGTGGTG |
| SSA2 R+1905 | AATACAGAGGAAAGCAAAAGTAAAAC |
| TMA10 F+159 | ACGAAGCAAAGTCTAACCCAAAG |
| TMA10 R+159 | TTTCATTGTTTTGGGAGTTAGAGC |
| UBI4 F+524 | GTAAGCAGCTAGAAGATGGTAGAACC |
| UBI4 R+524 | TGAATTTTCGACTTAACGTTGTCG |
| SCR1 F | CTCCACCTTCACCGCTGTTAG |
| SCR1 R | AAATATGGTTCAGGACACACTCC |

**Table S6. Primer used in construction of strains**

| Primer Name | Sequence (5' → 3') |
|---|---|
| **Primers used in construction of strains: SCY005** | |
| Fw Hsf1-mCherry-hphMX6 | AGGACCCGACAGAGTACAACGATCACCGCCTGCCCAAACGAGCTAAGAAA CCAGCTGAAGCTTCGTAC |
| Rv Hsf1-mCherry-hphMX6 | ACTATATTAAATGATTATATACGCTATTTAATGACCTTGCCCTGTGTA CCGCATAGGCCACTAGTG |
| **Primers used in construction of strains: SCY001, SCY003, SCY006, SCY012, SCY014, SCY016** | |
| Fw Med15-mCherry-hphMX6 | GTTCAGAACAATTCAATGTATGGGATTGGAATAATTGGACAAGTGCTACT CCAGCTGAAGCTTCGTAC |
| Rv Med15-mCherry-hphMX6 | CAAACGAAGTAACTTCAAAAGTATCAAAAGTATGGAAACTTCAAATGT CCGCATAGGCCACTAGTG |
| **Primers used in construction of strains: SCY002, SCY004, SCY007, SCY013, SCY015, SCY017** | |
| Fw Rpb3-mCherry-hphMX6 | ATGCATCTCAAATGGGTAATACTGGATCAGGAGGGTATGATAATGCTTGG CCAGCTGAAGCTTCGTAC |
| Rv Rpb3-mCherry-hphMX6 | TTCGGTTCGTTCACTTGTTTTTTTTCCTCTATTACGCCCACTTGAGAA CCGCATAGGCCACTAGTG |
| **Primers used in construction of strains: SCY008 and SCY009** | |
| Fw MCP-mCherry-Ura3 | ATAAATCATGTCGAAAGCTACATATAAGG |
| Rv MCP-mCherry-Ura3 | GCTCTAATTTGTGAGTTTAGTATACATGCATTTACTTATAATACAGTTTTTTA TTTTTTGCTTTTTCTCTTGAGGTCACATGATCG |
| Fw HSP12-MS2-loxp-KanMX6-loxp | TATGTTTCCGGTCGTGTCCACGGTGAAGAAGACCCAACCAAGAAGTAA GCCGCTCTAGAACTAGTGGATCC |
| Rv HSP12-MS2-loxp-KanMX6-loxp | ACACATCATAAAGAAAAAACCATGTAACTACAAAGAGTTCCGAAAGAT GCATAGGCCACTAGTGGATCTG |
| **Primers used in construction of strains: ASK213 and ASK214** | |
| Fw Med15-myc9 | GATTCTCCTGATGACCCATTCATGAC |
| Rv Med15-myc9 | CTGATGATAGTCAAGTCCATTGG |

| | |
|---|---|
| **Primers used in construction of strains: ASK215, ASK216 and ASK217** | |
| Fw Med15-myc13 | GTTCAGAACAATTCAATGTATGGGATTGGAATAA TTGGACAAGTGCTACTCG-<br>GATCCCCGGGTTAATTAACG |
| Rv Med15-myc13 | CACCAAACGAAGTAACTTCAAAAGTATCAAAAGT<br>ATGGAAACTTCAAATGTGAATTCGAGCTCGTTTAAAC |
| **Primers used in construction of strains: DPY1805 and DPY1818** | |
| Fw subCE2-AAA | CCAATAAGGCCCTATAAACAAAGAGCTGCTGCTAAAAATAGAGCCAAT-<br>TCCTCG |
| Rv subCE2-AAA | CGAGGAATTGGCTCTATTTTTAGCAGCAGCTCTTTGTTTATAGGGCCT-<br>TATTGG |
